# Adverse caregiving in infancy blunts neural processing of the mother: Translating across species

**DOI:** 10.1101/870261

**Authors:** Maya Opendak, Emma Theisen, Anna Blomkvist, Kaitlin Hollis, Teresa Lind, Emma Sarro, Johan N. Lundström, Nim Tottenham, Mary Dozier, Donald A Wilson, Regina M Sullivan

## Abstract

The roots of psychopathology frequently take shape during infancy in the context of parent-infant interactions and adversity. Yet, neurobiological mechanisms linking these processes during infancy remain elusive. Here, using responses to attachment figures among infants who experienced adversity as a benchmark, we assessed rat pup cortical Local Field Potentials (LFP) and behaviors exposed to adversity in response to maternal rough and nurturing handling by examining its impact on pup separation-reunion with the mother. We show that during adversity, pup cortical LFP dynamic range decreased during nurturing maternal behaviors, but was minimally impacted by rough handling. During reunion, adversity-experiencing pups showed aberrant interactions with mother and blunted cortical LFP. Blocking pup stress hormone during either adversity or reunion restored typical behavior, LFP power, and cross-frequency coupling. This translational approach suggests adversity-rearing produces a stress-induced aberrant neurobehavioral processing of the mother, which can be used as an early biomarker of later-life pathology.

## Introduction

A defining characteristic of young mammals is their obligatory attachment to their primary caregiver^1^. Despite this obligation, there are individual differences in the quality of this attachment which reflect the nature of caregiving received early in life and the way in which the infant psychologically represents routinized interactions with the parent. These individual differences are best identified by observation of brief separations followed by reunions between parent and infant. At young ages, separations from the parent are stressful, and therefore the way that the infant uses the parent for comfort at the time of reunion has proven diagnostic for classifying infants’ attachment quality within the range of typically developing children (i.e. secure or insecure) vs. attachment quality associated with later-life pathology (i.e. disorganized)^2,3^. The stress of the separation-reunion procedure is thought to be critical for uncovering these aberrant attachment styles as the roots of later-life socio-emotional difficulties, including poor stress-management, reactive attachment disorder, and future psychopathology. However, the causal and mechanistic pathways linking poor caregiving, attachment quality, and altered socio-emotional development have not been established.

Because the extant literature has relied on humans, current knowledge is based on correlational designs and investigative approaches that preclude invasive biological procedures. The current study aimed to overcome this methodological obstacle by assessing human and rodent reunion behaviors in parallel, and then within the causal rodent model, investigating the neurohormonal mechanisms of the observed attachment behaviors. Because the drive to reunite following separation is conserved across mammals, reunion behaviors can readily translate from humans to rodents to investigate neurohormonal mechanisms.

Here, we capitalize on the power of using animal models to understand human behavior by measuring localized brain activity and behavior during experimental treatment/testing, randomizing assignment, and assessing causation using a clinically-informed question. Although the pathways of attachment are complex, here we focus on the causal pathways between stress physiology and cortical function induced by the parent as a function of attachment quality. In particular, we focus on cortical oscillations - rhythmic neural activity that synchronizes the brain’s activity to coordinate functions within and across neuronal networks. Cortical oscillations are a biological cornerstone of brain development and are strongly governed by fluctuations in corticosterone^4,5^ and associated environmental stressors. While social stimuli are salient modifiers of neural oscillations throughout human development^6,7^, the attachment figure (biological or adoptive caregiver) is a particularly effective stimulus^8,9^.

We began our experimental approach using a translational framework to explore a causal link between attachment quality, infant experience with maternal care quality, and maternal regulation of the infant brain during separation-reunion (see Figure 1). Our testing of the infant’s response to reunion with the mother takes place within the framework of the Strange Situation Procedure (SSP). This procedure uses separation-induced distress followed by reunion with the parent and has diagnostic value for classifying infants’ attachment quality^2,10^. We present data in human infants who are at high-risk for maltreatment, which is paired with a rodent SSP (rSSP), where rat pups have been randomly assigned to adversity rearing with a maltreating mother or control rearing to assess attachment behaviors (see Figure 2 for SSP schematic). Further capitalizing on the rodent model, we probed direct assessment of causality by blocking pup stress hormone synthesis (metyrapone) for rescue during the rSSP. Next, again capitalizing on the rodent model, we probed the antecedents of rSSP deficits by measuring infant pups’ neurobehavioral responses during adversity rearing, thereby investigating which processes of the infant were perturbed by which behaviors of the mother in the maltreating context. Again, we probed direct assessment of causality in the rodent by blocking pup stress hormone synthesis (metyrapone) to prevent the behavioral and neurobiological aberrations identified in the rSSP.

**Figure 1.**
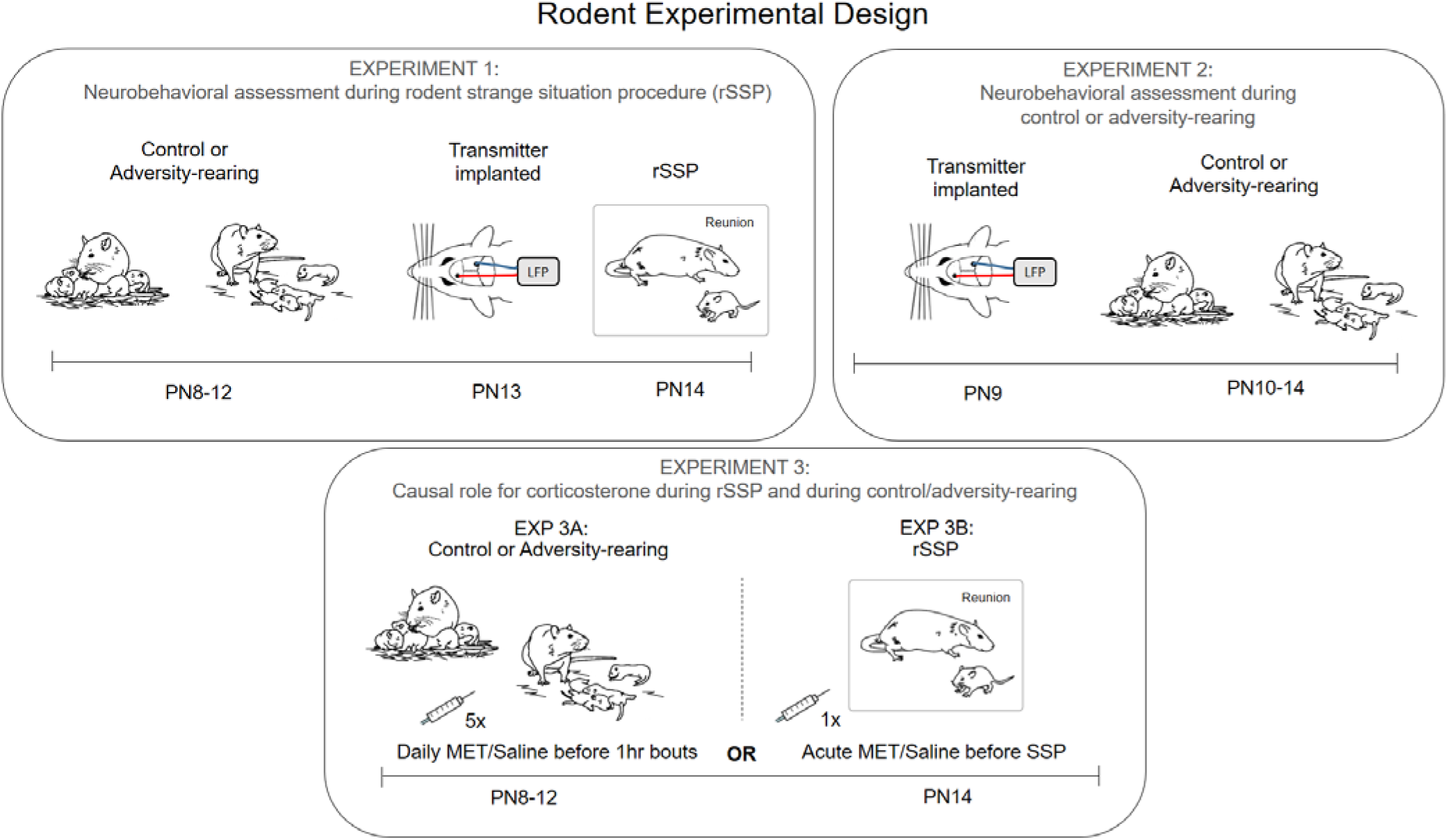
Rodent experimental design. In all rodent experiments, we used the Scarcity-Adversity model of low bedding, which induced maltreatment of pups in the rat mothers. In Experiment 1, adversity-rearing occurred from PN8-12 and at PN13-14, pups were removed from the nest and given the rodent rSSP while cortical LFP was recorded. In Experiment 2, neurobehavioral assessment of pups occurred during the control or adversity-rearing. Experiment 3 assessed causation in Experiment 1 and Experiment 2 results by blocking corticosterone synthesis. All pups experienced control or adversity-rearing from PN8-12, and then one cohort of pups received daily injections of metyrapone or saline 90 min before bedding was removed (5 administrations total, EXP 3A) while another cohort received a single injection 90 min before the rSSP procedure (EXP 3B).

**Figure 2.**
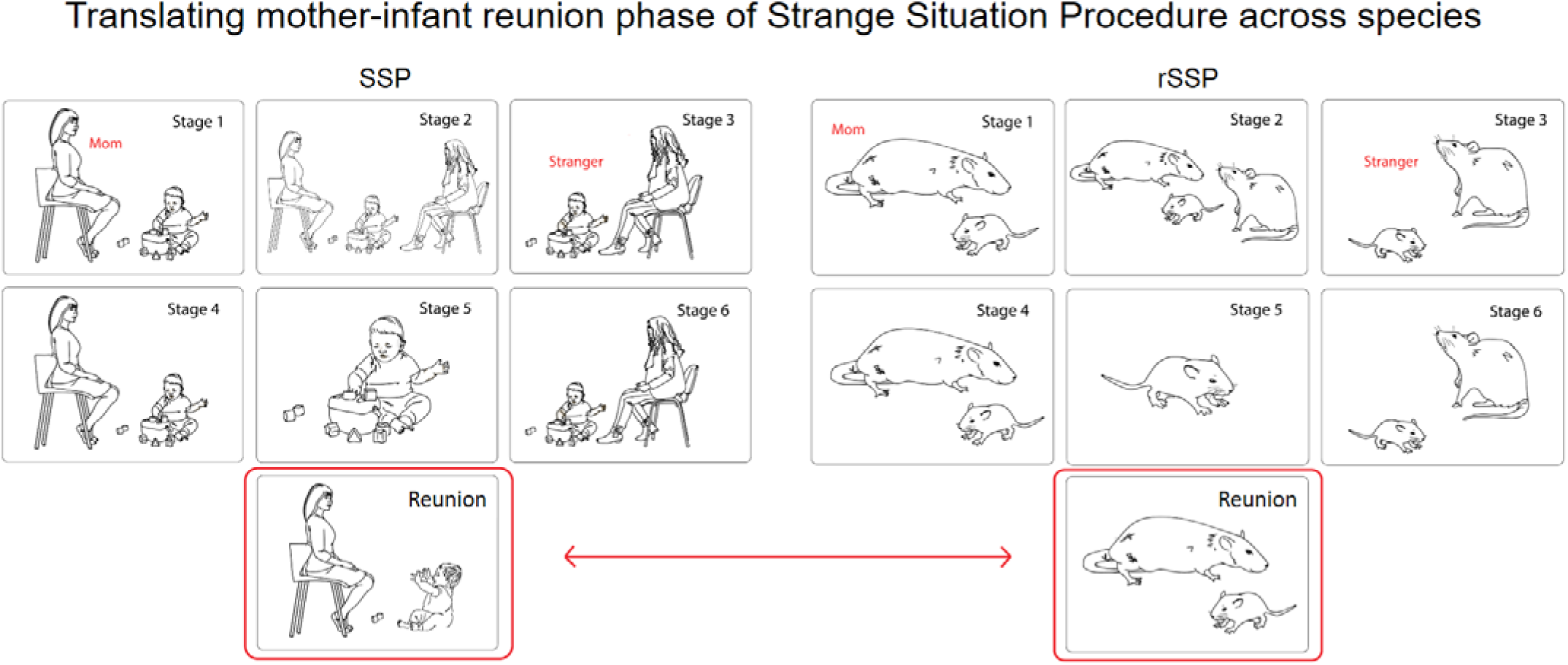
Translating mother-infant reunion phase of the Strange Situation Procedure across species. The SSP uncovers deficits in attachment behavior in children through a series of exposures to a stranger and separation-reunion with the mother, culminating in a final epoch of the child’s reunion with the mother^2,10^. Child behaviors during this reunion can signify the quality of attachment to the caregiver. While minor variations in attachment quality suggest individual differences in mother-infant interactions, attachment categories such as disorganized attachment highlight disruptive mother-infant attachment quality and predict later-life pathology^2,11^. Adaptation of the SSP to rodents (rSSP) uses a similar series of epochs with the mother. In rodents, where infants rely on the maternal odor for mother-infant interactions, the mother possessed the natural diet-dependent, learned maternal odor. A “stranger” was produced by feeding another lactating dam a different diet (Tekland) that suppressed the natural maternal odor and provided this mother with a novel maternal odor, which was unfamiliar to test pups. Importantly, pups reared with the “stranger” mother show strong attachment behaviors to this mother, while our test pups unfamiliar with this “stranger’s” maternal odor do not. Only the final reunion data from the SSP and rSSP is used to determine attachment quality. The SSP child subjects were 11-28 months old, while rat pups were PN13-14, an age range across species associated with some mobility and complete dependence on the caregiver for survival^12^.

## Methods and Materials

For detailed methods, please see Supplementary Materials.

### Human Subjects

Parents and their children were referred to the study because of risks for maltreatment, such as unstable living conditions, having mental health or substance abuse problems, or evidence of prior maltreatment. From this larger sample, two smaller samples were selected: a high-risk group that had at least 6 total risk factors and a control group that had no more than 5 or fewer total risk factors. Please see Tables S3 and S4 for additional information regarding human participants. All procedures were approved by the University of Delaware Institutional Review Board.

### Strange Situation Procedure in Children (SSP)

The SSP is designed to permit assessment of children’s reliance on the parent when they are distressed and involves a standard series of separations and reunions of the child with the parent^10^. Attachment assessment occurs primarily during the two reunions with the parent; child proximity seeking, contact maintenance, avoidance, and resistance are coded during reunions, and disorganized behavior is coded throughout. Please see Supplementary Methods and Table S5 for additional information on how child behaviors were coded during this procedure.

### Rodent Subjects

Rat pups were bred and housed with their mother in standard cages with wood chips and *ad libitum* food and water in an environmentally-controlled room. All procedures pertaining to the use of live rats were approved by the Nathan Kline Institute Institutional Animal Care and Use Committee and followed National Institutes of Health guidelines.

### Scarcity-Adversity Modeling: Infant maltreatment in rodents

Our Scarcity-Adversity model of low bedding (adversity-reared) occurs in the home cage with a solid floor and robustly increases maternal rough treatment of pups^13,14^. Pups live in either typical nest environments (control) or with adversity-rearing mothers given insufficient bedding for nest building (100ml vs. 4500ml, both cleaned 2-3X). This manipulation decreases the mother’s ability to construct a nest, resulting in frequent nest building and transporting/rough handling of pups, although pups exhibit normal weight gain after five continuous days of adversity-rearing^15^. In the present study, we use either a chronic [5 days, postnatal days (PN8-12)] treatment (Experiment 1, rSSP) or, in a separate cohort of animals, brief 1 hr bouts of adversity-rearing twice a day (Experiment 2, during adversity-rearing) between the pup ages of PN10-14.

### Strange Situation Procedure in Rodents (rSSP)

The rodent SSP (rSSP) was conducted after the control or adversity-rearing, using the mother and a “stranger” rat mother (novel maternal odor). In the first two weeks of life, rat pups primarily identify their mother by the maternal odor, which is learned^16,17^ as hearing and vision senses are functionally immature^18,19^. With the mother and stranger anesthetized, the rat pup was placed in a small plastic chamber for the entire test with mother and stranger presence manipulated (epochs summarized in Figure 2). Similar to humans, the final reunion epoch was used for data.

### In-nest observations during and after scarcity-adversity procedure

Videos were recorded and hand-scored by highly-trained raters to validate that the adversity-rearing treatment induced maltreatment by mothers (Table 1) and post adversity-rearing maternal care was back to baseline (Fig. S2). In SSP experiments 1 and 3, raters were blinded to the previous rearing condition of pups. In Experiment 2, during LFP recordings, home cage videos were taken during the adversity-rearing to define nurturing and maltreatment maternal behavior.

### Surgical procedures

One to two days prior to an experiment, pups were anesthetized (isoflurane), implanted with a stainless steel electrode targeting the neocortex (~2 mm anterior to bregma, ~2mm laterally over the left hemisphere, ~1.0mm ventral to brain surface), and connected to a telemetry pack (ETA-F10 telemetric device, DSI) inserted subcutaneously on the back^20^. Following recovery from anesthesia (<30min) pups were returned to the mother, littermates, and nest.

### LFP recordings

The telemetry neural signals were filtered (0.5 to 200 Hz), digitized at 2 kHz with Spike2 software (CED, Inc.), and analyzed offline. During the rSSP, we recorded spontaneous LFP that was analyzed by epoch and across pup behaviors toward the mother and stranger. LFP was also recorded during control and adversity-rearing for 4-5 consecutive days in the freely behaving infants interacting with the mother during daily hour-long nesting bouts. LFP sessions were videotaped and behaviors noted on the neural trace for off-line analysis^20^.

### Data analysis

Fast Fourier Transform (FFT) power analyses were performed on raw LFP data in intervals taken from sections of daily neural traces that correlate with specific nurturing and maltreating behaviors to quantify LFP oscillatory power in 2.9 Hz frequency bins from 0–100 Hz (Hanning)^20^. Power in the delta (0-5 Hz), theta (5–15 Hz), beta (15-35 Hz), and gamma (35-100) Hz) frequency bands was calculated for each specified window.

### Cross-frequency coupling analysis

Data were analyzed as previously described^21^ and in Supp. Methods. LFP traces were filtered into theta (1.5–12 Hz) and gamma (35–80 Hz). Briefly, cycles (the nadir of negative troughs) were then detected based on threshold crossing from filtered signals. Phase locking was assessed for each oscillatory frequency band as follows: Experiment 1 assessments during SSP epoch 7 (5 min); Experiment 2 assessments during 1 hour bouts of control and adversity-rearing^22^.

### Corticosterone manipulation

In Experiment 3, separate cohorts of pups received intraperitoneal injections of the corticosterone inhibitor metyrapone HCL (50 mg/kg, Sigma) or an equal volume of 0.9% saline. Drug administration occurred 90 mins before either the daily 1 hr bout during control or adversity-rearing or before the rSSP.

### Statistical analysis

Depending on the number of groups to be analyzed, behavioral and LFP data were analyzed with Student’s t-tests and one-or two-way analysis of variance (ANOVA), followed by post-hoc Bonferroni tests, or X^2^ analysis and circular statistics (coupling data). Data used for figures are expressed as mean (±SEM) and in all cases, differences were considered significant when *p*<0.05.

## Results

### Experiment 1: Across species, compromised early care disrupts attachment quality reflected in separation and reunion with the parent

#### Maltreatment risk and the Strange Situation Procedure (SSP) in humans

This procedure uses separation-induced distress followed by reunion with the parent to characterize how the infant uses the parent for comfort, which has diagnostic value for classifying infant’s attachment as secure (i.e. low risk for later psychopathology) vs. disorganized (i.e. high risk for later psychopathology)^2,10^. Our SSP subjects were children [N=21; 12.1-28.3 months of age (*M*=19.9, *SD*=5.6)] and their mothers separated into two groups: a high-risk group that had at least 6 risk factors (7m4f) and a low-risk group that had no more than 5 risk factors (6m4f). The relationship between risk for maltreatment and attachment classification was significant [Fig. 3A; X^2^(1,21)=3.834, *p*=0.05; relative risk=2.867)]. Specifically, infants with fewer risk factors were more likely to show secure, organized attachment, such as approaching the parent and being calmed by the parent’s return whereas infants with more risk factors were more likely to exhibit disorganized attachment behaviors [X^2^(1,21)=14.32, *p*=0.0002, relative risk=5.5], such as initial greeting followed by avoidance or crying at a distance from the mother. Parents from the high-risk group were less sensitive to their infants than low-risk parents (*r* = −0.42, *p* = 0.008), and more sensitive parenting was associated with fewer disorganized behaviors than less sensitive parenting (*r* =-.44, *p =* 0.047) (Fig. S1). Thus, these data replicate a common finding in the literature^11,23^, where risk for maltreatment was associated with a greater risk for disorganized attachment behaviors.

**Figure 3.**
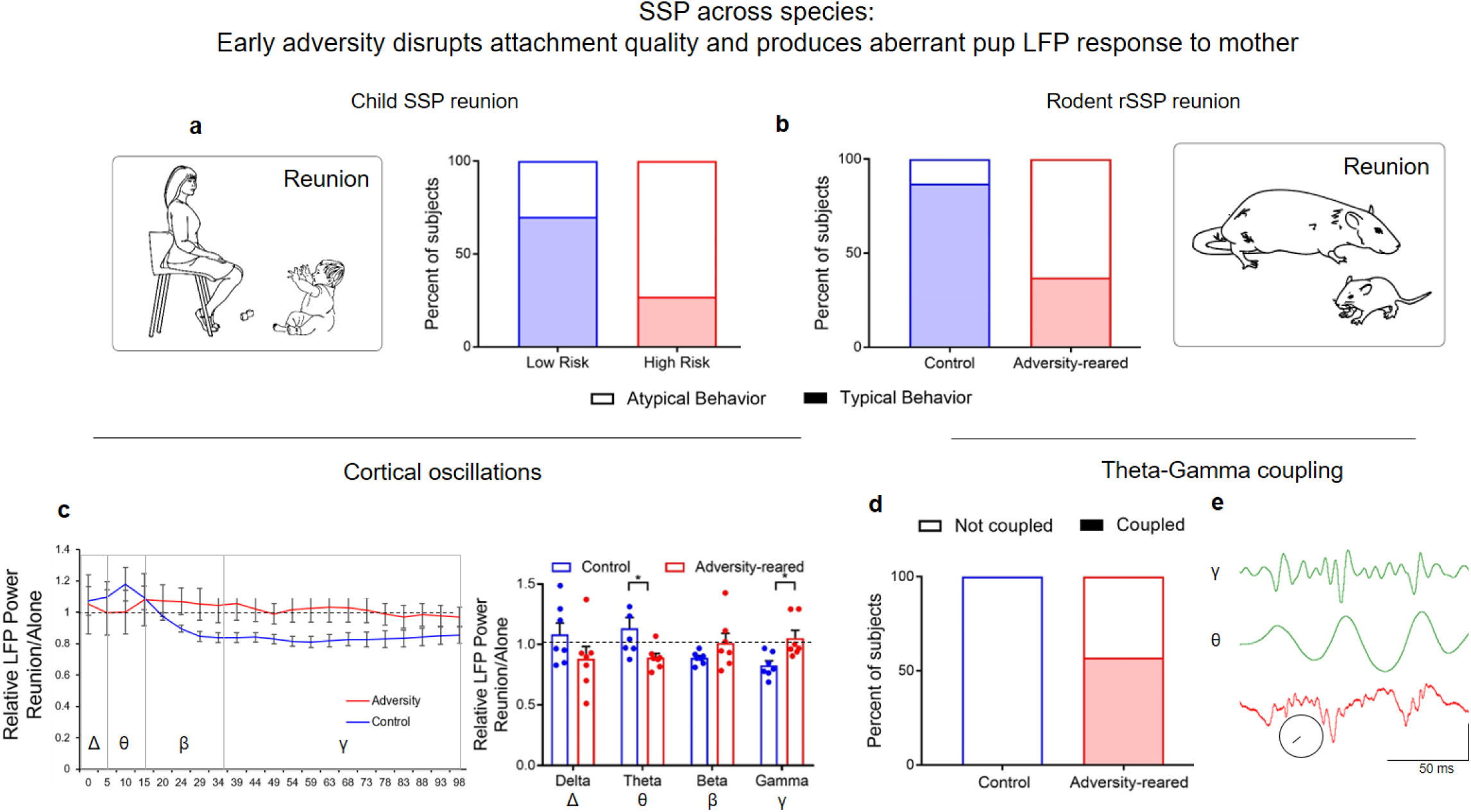
Across species, compromised early care disrupts attachment uncovered by separation and reunion with the caregiver in the SSP. A) In children, biographical factors (eg. SES, housing stability) generated classifications of risk. During the SSP, prosocial behaviors were observed more often in low risk children, while atypical reunion behaviors were more frequently observed in high risk children. B) Adversity-reared rat pups showed atypical responses to the caregiver during the SSP reunion epoch. Across species, atypical reunion behaviors observed in the child SSP and rodent rSSP included contradictory responses to the parent or misdirected behaviors. C) In rat pups, reunion with the mother decreased high frequency oscillations in the control-reared pups (normalized to power during epoch 5: pup alone, data from 5 control and 5 adversity-reared litters). This decrease was not observed in maltreated pups; unlike controls, theta power was decreased. Bars represent mean ± SEM of power collapsed into frequency bands. Dashed line = no change from pup alone. D) In rat pups, Theta-gamma coupling was observed more reliably in maltreated pups than control-reared pups. E) Rat pup example of raw voltage trace (red) and gamma/theta filters (green); scale bar: vertical is 10 microvolts in theta/gamma, 25 microvolts in raw trace. Circle shows phase locking vector length = 0.0581, - 122.36°. *p<0.05. Source data are provided as a Source Data file.

There are several remaining unanswered questions in the human studies. For one, is the association between risk for maltreatment and attachment behaviors causal or simply correlational? Second, what is it about the parental behavior that changes the infant’s attachment relationship? And third, what are the neurobiological underpinnings in the maltreated infant’s developing brain that are giving rise to these disorganized attachment behaviors, and are they reversible? To answer these questions, we developed a rodent Strange Situation Procedure (rSSP).

**Table 1.**
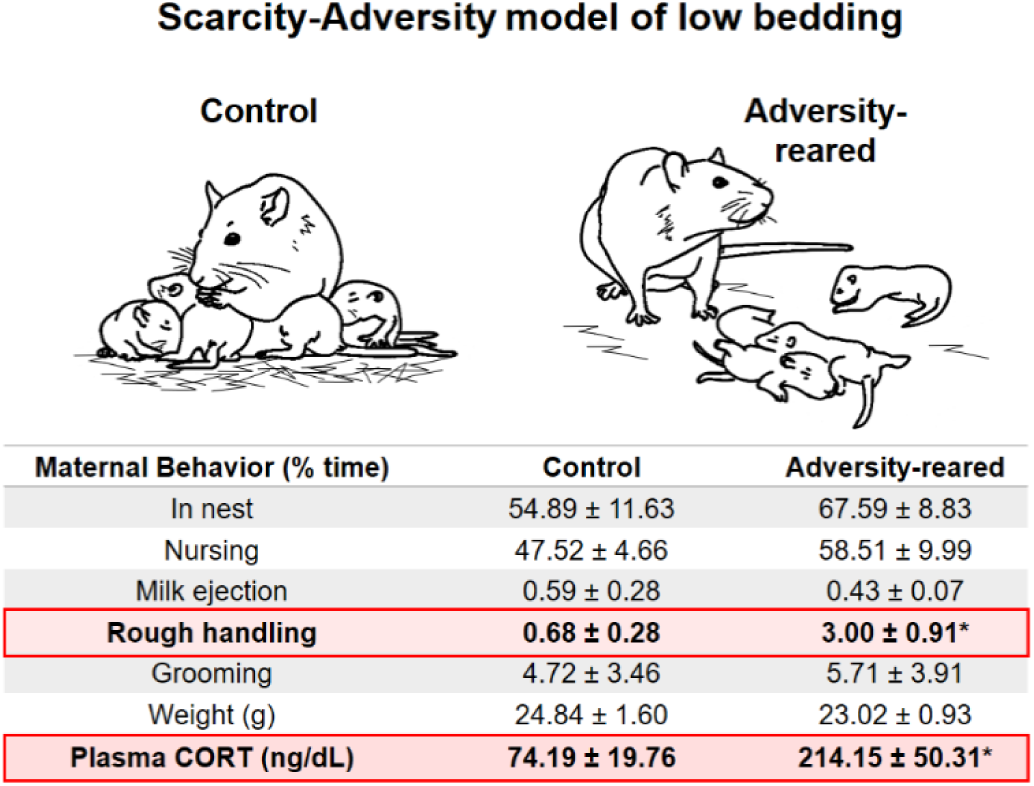
Relative occurrence of maternal behaviors within adversity-reared vs. control-reared litters. Decreasing bedding materials from 4000ml to 100ml produces robust and immediate changes in maternal behavior. Source data are provided as a Source Data file. The relative frequency of rough handling (dragging and stepping on pups) is more frequent in the maltreating mothers, behaviors which typically occur as the mother enters and leaves the nest. Pup plasma corticosterone levels are elevated after 5 days of this adversity-rearing. This procedure is documented to induce later life neurobehavioral psychopathologies, which are summarized with citations in Table S1.

|  | Control | Adversity-reared |
| --- | --- | --- |
| Maternal Behavior (% time) | Control | Adversity-reared |
| In nest | 54.89 ± 11.63 | 67.59 ± 8.83 |
| Nursing | 47.52 ± 4.66 | 58.51 ± 9.99 |
| Milk ejection | 0.59 ± 0.28 | 0.43 ± 0.07 |
| **Rough handling** | **0.68 ± 0.28** | **3.00 ± 0.91\*** |
| Grooming | 4.72 ± 3.46 | 5.71 ± 3.91 |
| Weight (g) | 24.84 ± 1.60 | 23.02 ± 0.93 |
| **Plasma CORT (ng/dL)** | **74.19 ± 19.76** | **214.15 ± 50.31\*** |

#### Maltreatment and the Strange Situation Procedure (rSSP) in rodents

Across species, attachment is preserved when infants experience repeated maltreatment by the caregiver, although the human literature shows the resulting attachment quality is aberrant ^2,24^. Here, we capitalized on this phylogenetically preserved system and induced maternal maltreatment of rat pups to question whether it also disrupts rodent infant attachment. From PN8-PN12, we randomly assigned half (n=8) of the pups to the maltreatment-inducing Scarcity-Adversity paradigm, while the other half (n=8) were reared in control housing. Here and in all experiments, samples included approximately equal males and females (see Supp. Methods for specific distribution information). The relative frequency of rough treatment of pups by the Scarcity-Adversity mothers were significantly higher than control mothers (Table 1, *t*(4)=2.98, *p*=0.027), while nurturing behaviors (nursing, grooming) occurred at control levels (all p’s>0.05), consistent with previous research^15,25^.

With two groups of rat pups, those with maltreatment histories and those without, we proceeded to the rSSP (PN13-14) (Experiment 1). We, a research team with rodent and human expertise in attachment, developed the rSSP to accommodate infant rodents and simulate the human procedure. Such translation is made possible by the fact that the human SSP was originally designed to be ecologically relevant to the young altricial infant who is mobile but still depends on the parent for soothing check-ins if stressed or anxious. This developmental ecology is maintained across the human, nonhuman primates, and rodent infant^12^. In the rSSP, as in the human SSP, 7 epochs are comprised of separations, reunions, and introductions of a “stranger” rat mother, produced by changing the maternal odor via the diet (see Supp. Methods; epochs summarized in Figure 2). Similar to humans, the final epoch (“Reunion”) was used for data.

Adversity-reared rodents showed a behavior pattern during the rSSP that was similar to that observed in high-risk human infants in the SSP. For attachment behaviors upon final reunion with the mother rat, adversity had a significant effect. Pups from control mothers were more likely to show typical attachment behaviors upon reunion, such as approaching the mother, nursing, and sleeping at the mother’s ventrum [X^2^(1,16)=2.618, *p*=0.05; relative risk=1.5]. In contrast, pups that were maltreated induced by the Scarcity-Adversity model of low bedding (“Adversity” in graphs) were less likely to show these behaviors, instead exhibiting atypical behaviors such as sleeping behind the mother’s back and sleeping alone [Fig. 3B, X^2^(1,16)=4.267, *p*=0.039, relative risk=0.2]. Importantly, nursing behavior in the nest did not differ between groups, suggesting the rSSP testing procedure uncovered group differences (Table 1), similar to the SSP in children. As shown in Figure 3A-B, behaviors exhibited by adversity-reared rat pups paralleled those exhibited by the children with high maltreatment risk.

This behavioral parallel is important because it demonstrates a shared developmental ecology between altricial young humans and rats that depends on forming robust attachment relationships with their parents regardless of the quality of maternal care, although maltreatment within attachment produces suboptimal attachment quality. Additionally, these data provide empirical evidence that the aberrant behaviors exhibited by infant rats in the rSSP are caused by the maternal behaviors (and are not merely correlational) through a randomization process that can rarely be executed in humans.

#### LFP cortical response to reunion with mother is blunted in adversity-reared rat pups

This behavioral parallel provides the platform for interrogation of the mechanisms that give rise to SSP behaviors that are changed by maltreatment. Next, we assessed whether pups’ brain responses (i.e., LFPs) to maltreating mothers were altered during the rSSP. Maternal regulation of infant brain oscillations has received minimal attention in humans, although a recent study has shown that in older children (~11 years old), maternal cues presented during magnetoencephalography (MEG) recording enhanced theta and gamma frequency band activity^9^. Importantly, in mother-child pairs associated with conflict style, the presence of the mother failed to produce as robust a change in the infant’s EEG gamma oscillations compared to that seen in well-adjusted, mother-infant dyads^8^. Together, this work suggests that neural oscillations are significantly influenced by the attachment figure but potentially compromised by poor attachment. During the last epoch of the rSSP, when the pup was reunited with the mother, control-reared pups showed a decrease in LFP gamma band power compared to the epoch when they were alone (Fig. 3C). This corroborates previous research showing that maternal presence in the nest decreases cortical LFP gamma band power compared to when pups are alone^20^. However, adversity-reared pups failed to exhibit a similar decrease in gamma power during reunion with the mother. ANOVA showed a significant interaction between rearing condition (control vs. adversity-reared) and frequency band (delta, theta, beta, gamma)[*F*(3,48)=5.415, *p*=0.003; no main effect of frequency band (*F*(3,48)=0.439, *p*=0.727) or rearing condition (*F*(1,48)=0.229, *p*=0.635)]. *Post hoc* comparisons showed that the difference between control and adversity-reared pups was in both theta and gamma frequency bands (theta: control = 1.135±0.088, adversity-reared = 0.892 ±0 .036, *t*(48)=2.424, *p*=0.019; gamma: control = 0.827 ± 0.038, adversity-reared = 1.053 ± 0.064, *t*(48)=2.251, *p*=0.029).

Next, we assessed theta-gamma coupling during the reunion with the mother (Fig. 3D). Cross-frequency coupling allows assessment of how effectively circuits can regulate local and network communication at different spatiotemporal scales and has been shown to be sensitive to developmental events^5,26,27^. Briefly, LFP recordings during the reunion epoch (5 min) were filtered into theta and gamma bands and cycles (troughs) were detected. To avoid confound with differences in oscillation amplitudes, average number of gamma events per theta event was balanced between conditions (control = 44.85 ± 5.135, adversity-reared = 40.3 ± 5.928, *t*(12)=0.578, *p*=0.577). We observed more reliable theta-gamma coupling in pups that had been maltreated than those that had not been maltreated (X^2^(1,14)=5.6, *p*=0.018, relative risk=8). Whereas theta-gamma coupling was observed in 4 of 7 adversity-reared pups, coupling was not observed in any of the pups reared in controls. Mean vector length doubled between conditions (control, 0.019 ± 0.002 vs. adversity-reared, 0.043 ± 0.011; *t*(12)=2.137, *p*=0.025).

### Experiment 2: During maltreatment, pups exhibit robust alterations in LFP response to the nurturing, not maltreating, behaviors of their mothers

Next, to better understand the antecedents of these observed neurobehavioral changes within the rSSP, we moved our behavioral and cortical oscillations assessment to within the nest during control and adversity-rearing treatment in the home cage. Pup and maternal behaviors were observed and LFPs telemetrically recorded during 1 hour bouts of Scarcity-Adversity model of low bedding (“Adversity-reared”, n=6) and control (n=7) across several days. Whereas pups in the rSSP interacted with an anesthetized mother, here we assessed the temporal course of untethered and experimentally undisturbed pup interaction with the mother in the home cage.

Pup cortical oscillations were analyzed during specific maternal behaviors towards pups: some maternal behaviors produced differences in oscillations between rearing conditions, while others did not (summarized in Figure 7). No differences were detected between control vs. adversity-reared pups when the mother was out of the nest and during non-nutritive sucking. [Fig 4A; Mother out of nest - two-way ANOVA, frequency band (delta, theta, beta, gamma) x rearing (control vs. adversity-reared), no significant interaction (*F*(3,44)=0.877, *p*=0.460, main effect of frequency (*F*(3,44)=4.753, *p*=0.006); and no main effect of rearing *F*(1,44)=1.742, *p*=0.194)] and [Fig 4B; Non-nutritive nursing - two-way ANOVA, frequency x rearing, no significant interaction (*F*(3,44)=0.351, *p*=0.789, main effect of frequency (*F*(3,44) = 3.504, *p*=0.023); and no main effect of rearing *F*(1,44)=0.277, *p*=0.601)]. Given that pups were homogeneously quiescent during non-nutritive nursing, individual LFP power during this nursing was used to normalize pup oscillations in response to specific maternal presence and behaviors.

**Figure 4.**
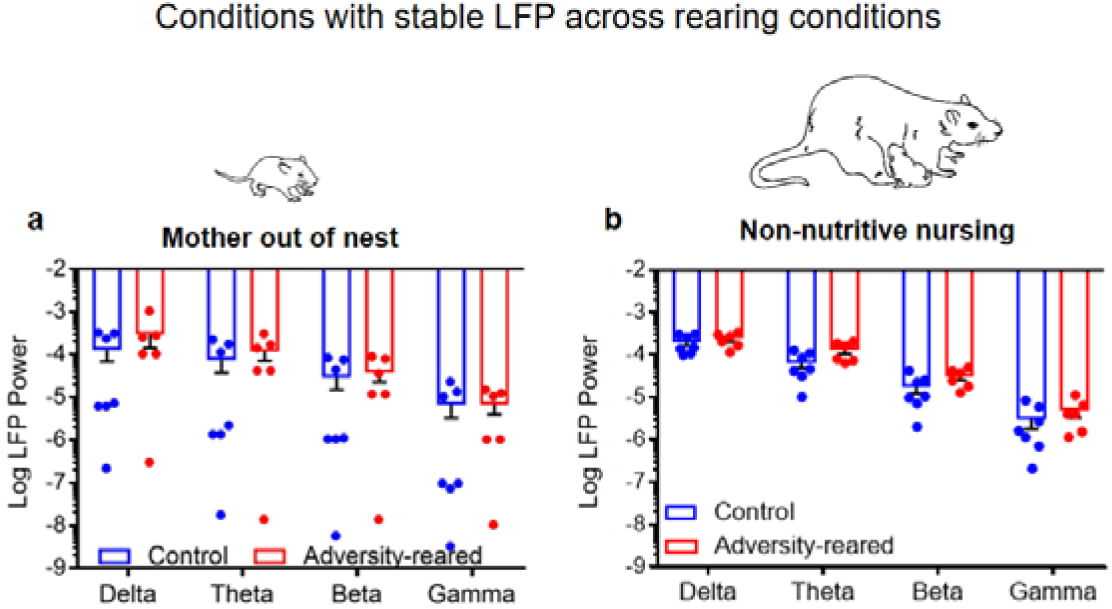
Stable cortical oscillations across rearing conditions during specific behavioral states. A) When the pup is alone/away from the mother, LFP power does not differ between rearing conditions. B) Cortical LFP power did not differ between conditions when pups are non-nutritively nursing. This measure (“baseline”) was used to normalize evoked responses to maternal inputs. Bars indicate mean ± SEM. Source data are provided as a Source Data file.

**Figure 7.**
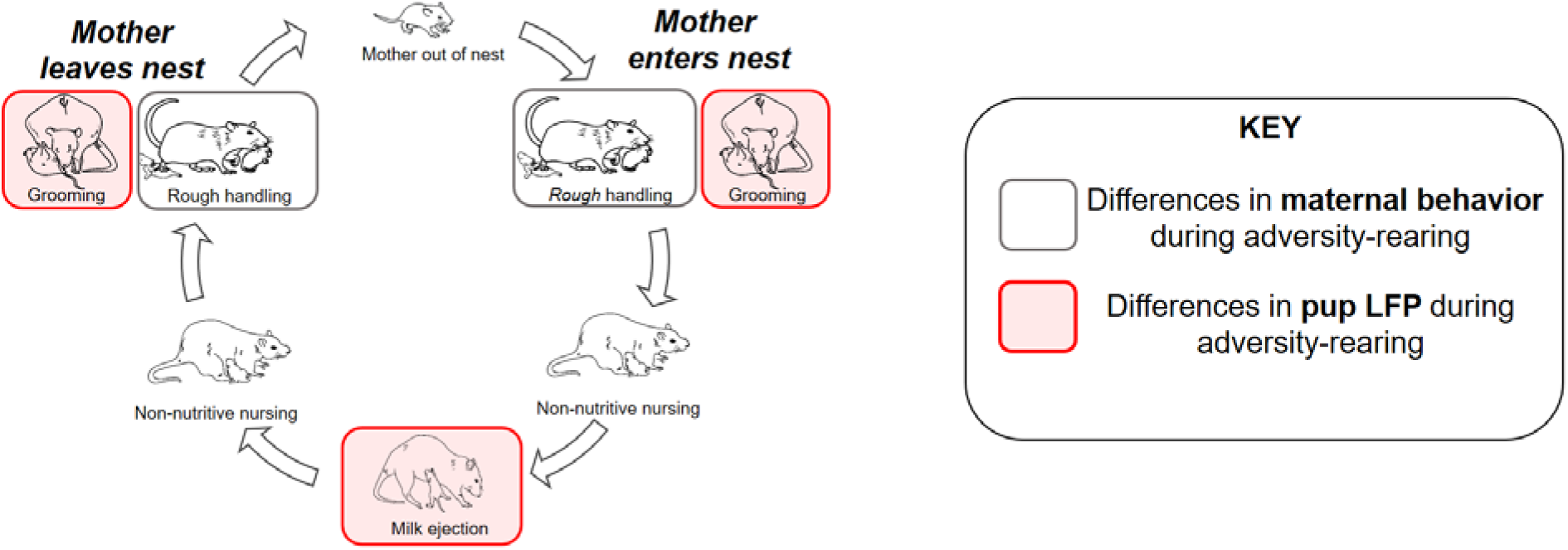
Differences in maternal behavior during maltreatment do not align with altered pup cortical processing of maternal behavior. The typical sequence of maternal behavior does not change between control and maltreatment rearing: Pups are left alone in the nest as the mother attends to her biological needs. Upon returning to the nest, the mother adjusts bedding, moves, and grooms pups as she begins to nurse. During most of nursing, pups are non-nutritively nursing, which is interrupted periodically by a milk ejection, which produces a robust stretch reflex in pups. Although low bedding induces increased rough handling by mothers (grey border) upon entering/exiting nest, specific maternal nurturing behaviors (red) were associated with the most robust differences in pups’ cortical reactivity between the control and low bedding groups. Please see Supp. Materials for expanded discussion of mother-pup interaction within nest.

Other maternal behaviors produced statistically different responses in control vs. adversity-reared pups. While the Scarcity-Adversity rearing increased rough handling (dragging/stepping on pups), the occasional occurrence of rough handling in control mothers allowed us to compare rough handling across rearing conditions. Surprisingly, during maternal rough handling of pups, LFP responses in adversity-reared and control pups only differed in the delta band [Fig. 5E-F; two-way ANOVA, frequency x rearing, no main effect of frequency (*F*(3,44)=2.592, *p*=0.065); main effect of rearing (*F*(1,44)=4.303, *p*=0.044); *post hoc* delta: control = 1.317 ± 0.32, adversity-reared = 0.673 ± 0.115, *t*(44)=2.51, *p*=0.016]. Delta frequency band activity is normally elevated during slow-wave sleep and reduced during other states. While it is unlikely that either control or adversity-treated pups were sleeping during rough handling, the significant reduction in delta band power in adversity-treated pups during rough handling may reflect greater arousal in these pups compared to controls during these adverse events. It should be noted that delta band power can be strongly impacted by movement artifacts and care was taken here and elsewhere to select artifact-free behavioral bouts.

**Figure 5.**
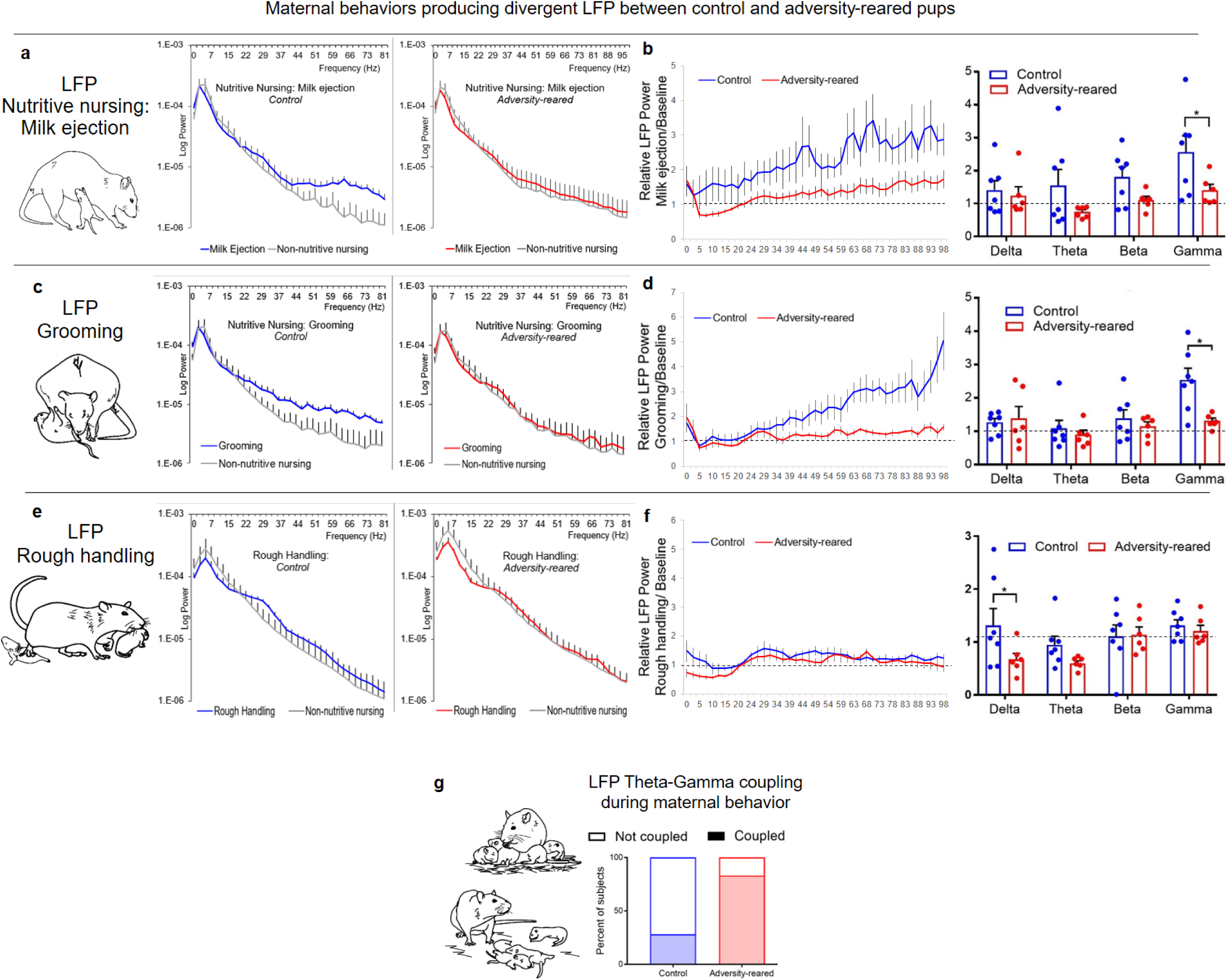
During adversity rearing, pup LFP response to specific maternal behaviors is decreased. A) While milk ejection increased pups’ oscillatory power in the theta and gamma ranges, these inputs failed to modulate oscillations in the same pups during maltreatment. Left, log LFP power; B) LFP power normalized to LFP during non-nutritive nursing (between milk ejections, “baseline”). C) Grooming failed to modulate pup cortical oscillations when this occurred within adversity rearing. Data normalized to baseline are shown in D. E) Specific oscillatory responses to maternal behavior during actual rough handling (i.e. during stepping on pup or dragging pups) differed minimally between the control and adversity-reared conditions, limited to the delta range, shown normalized in (F). G) Cortical theta-gamma coupling was more reliably observed during bouts of adversity rearing. *p<0.05. Error bars = SEM. Dashed line = no change from baseline (non-nutritive nursing LFP power). Source data are provided as a Source Data file.

However, the most robust LFP differences were detected between adversity-reared and control pups’ LFP oscillations during *nurturing* maternal behaviors. The pups in the control environment showed enhanced high frequency power to various maternal cues compared to LFP activity during non-nutritive nursing (baseline) conditions. Specifically, during both grooming and nutritive sucking (milk ejection), control pups showed an increase in high frequency LFP power, which was measured for ten-second periods after their initiation. During milk ejection in control-reared pups, we observed a surge in high frequency oscillations compared to non-nutritive nursing baseline (Fig. 5A-B, beta: *t*(6)=2.631, *p*=0.039; gamma: *t*(6)=3.159, *p*=0.019). Grooming similarly produced an increase in LFP power in the gamma range (*t*(6)=4.314, *p*=0.005), consistent with previous work in typically-reared pups^20^.

During adversity-rearing, the frequency of nurturing behaviors (nutritive/non-nutritive nursing and grooming) remained at control levels. However, in sharp contrast to control-reared pups, these same nurturing behaviors failed to modulate the adversity-reared pups’ cortical oscillations. Specifically, we observed significant effects of rearing condition (control vs. adversity-reared) on pup LFP power following milk ejection and grooming [Fig. 5C-D; milk ejection: two-way ANOVA, rearing x frequency, main effect of rearing (*F*(1,44)=9.102, *p*=0.004); grooming: two-way ANOVA, rearing x frequency, interaction between rearing and frequency (F(3,44)=3.029, *p*=0.039, main effect of frequency (F(3,44)=5.451, *p*=0.003), main effect of rearing (*F*(1,44)=5.205, *p*=0.027)]. *Post hoc* comparisons show a significant effect of Scarcity-Adversity treatment on gamma modulation by milk ejections (control = 2.551±0.284, adversity-reared = 1.4±0.269, *t*(44)=2.47, *p*=0.017). In addition, *post hoc* tests show that the Scarcity-Adversity treatment blunted modulatory effects of grooming in the gamma band (control = 2.523 ± 0.353, adversity-reared = 1.298 ± 0.083, *t*(44)=3.651, *p*<0.001).

Finally, we observed a significant change in cortical theta-gamma phase coupling (Fig. 5G; X^2^(1,13)=3.899, *p*=0.024, relative risk=2.94), with increased occurrence of coupling in pups during bouts of adversity rearing compared to control. These results parallel the pattern of more reliable coupling observed in adversity-reared pups in the rSSP. Mean phase coupling vector length doubled during adversity (control: 0.015±0.006, adversity-rearing: 0.036±0.015) though this did not reach statistical significance (*t*(11)=1.341, *p*=0.207). Average number of gamma events per theta event was balanced between conditions (*p*>0.05).

### Experiment 3: Lowering pup stress hormone corticosterone (CORT) levels restores maternal neurobehavioral regulation of adversity-reared pups

Increased stress hormones have long been associated with both initiating infant neurobehavioral pathology and its expression^23^, and animal and human research has shown stress hormones are associated with long-term outcomes of early life trauma^28^. For this reason, we next leveraged the power of our animal model to test if there is a causal link between stress hormone levels and the neurobehavioral deficits induced by Scarcity-Adversity treatment.

As shown in Table 1, plasma CORT levels increased in pups during adversity-rearing. We therefore administered the CORT synthesis inhibitor metyrapone (“Met”, Sigma, 50 mg/kg) or saline 90 minutes before daily bouts of adversity/control for 5 days. Importantly, daily injections were performed very rapidly without pup vocalizations and did not alter maternal rough handling of pups (Table S2). This treatment rescued typical attachment behavior towards the mother [Fig. 6A: two-way ANOVA, rearing (control vs. adversity) x drug (saline vs. metyrapone), significant interaction (*F*(1,20)=5.19, *p* =0.034); main effect of rearing (*F*(1,20)=8.061, *p*=0.01); main effect of drug (*F*(1,20)=5.48, *p*=0.029)]. *Post hoc* analyses showed that while saline-treated adversity-reared pups showed a significant increase in atypical attachment behaviors compared to saline-treated controls (305.667 ± 151.656 vs. 1211 ± 310.7, *t*(20)=3.266, *p*=0.004), attachment behavior did not differ between Met-treated, adversity-reared pups (208 ± 121.341) and saline controls (*t*(20)=0.352, *p*=0.728).

**Figure 6.**
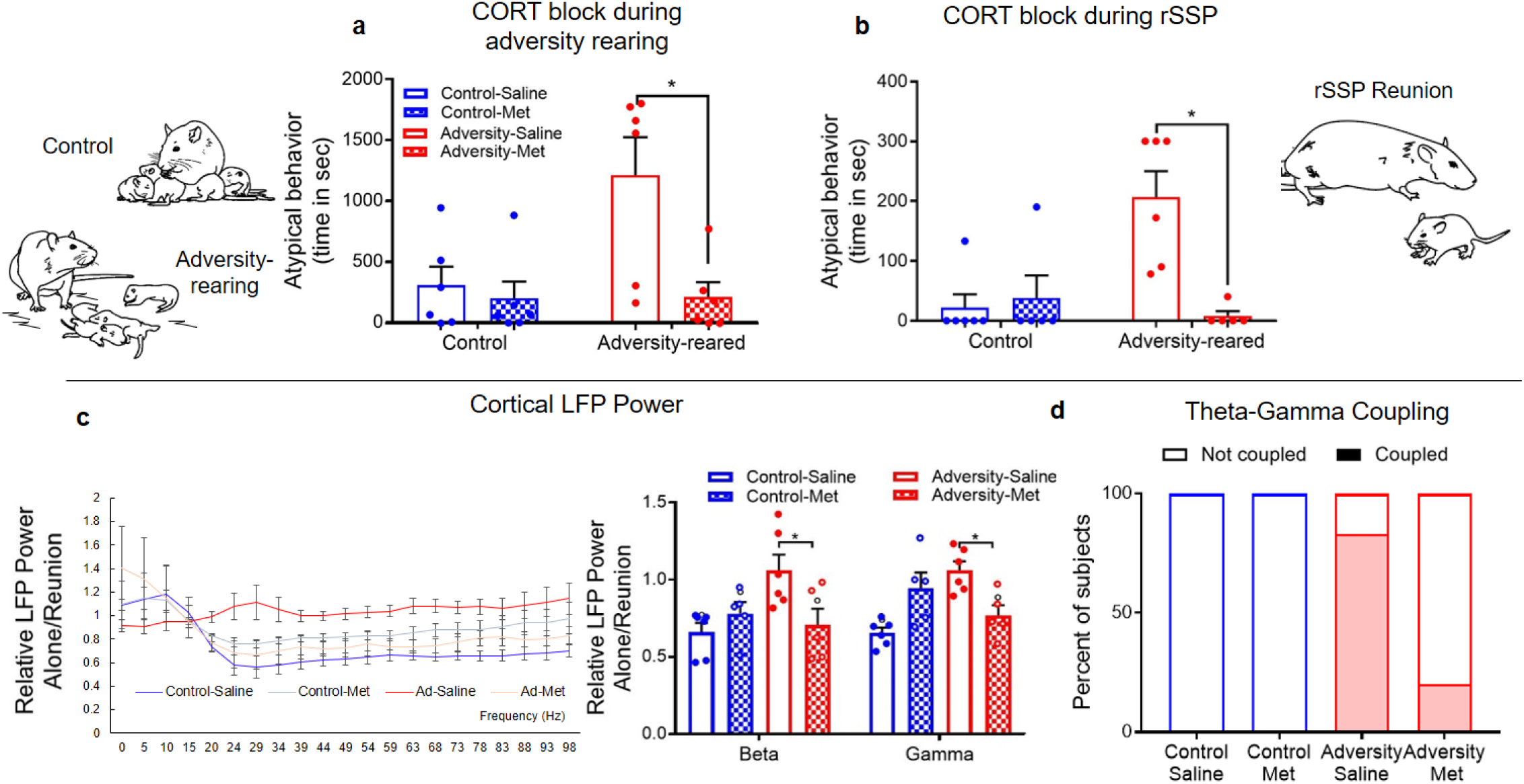
Decreasing the stress hormone corticosterone (CORT) rescues maternal regulation of infant attachment behavior and cortical oscillations. A) Blocking CORT synthesis during maltreatment prohibited expression of atypical behavior during a 30-minute social behavior test outside of the nest. B) Blocking CORT synthesis during SSP prevented expression of atypical attachment behavior during reunion with the mother. 90 minutes before SSP, pups received the CORT synthesis inhibitor metyrapone (50 mg/kg) or saline. C) In an internal replication of Exp. 1, saline-treated, adversity-reared pups failed to exhibit decreased beta and gamma oscillations upon reunion with the mother; this maternal regulation was restored if they were administered metyrapone before the test. Bars represent mean ± SEM of power collapsed into frequency bands. d) CORT inhibition in maltreated pups restored control-like coupling during reunion with the mother in the rSSP. Dashed line = no change from baseline. ⁰ significantly different from baseline (1). *p<0.05. Source data are provided as a Source Data file.

This same pattern of behavioral rescue was observed when metyrapone was administered 90 minutes before the rSSP [Fig. 6B; two-way ANOVA, rearing x drug, significant interaction (*F*(1,18)=11.03, *p*=0.004), main effects of drug (*F*(1,18)= 8.103, *p*=0.011) and rearing (*F*(1,18)=5.722, *p*=0.028)]. *Post hoc* analyses showed that while saline-treated, adversity-reared pups showed a significant increase in atypical attachment behaviors compared to saline-treated controls (Control Sal = 22.167 ± 22.167, Adversity Sal = 206.667 ± 43.778, *t*(18)=4.237, *p*=0.005), attachment behavior did not differ between adversity-reared pups treated with metyrapone and saline controls (*t*(18)=0.31, *p*=0.76).

These behavioral changes were also reflected in pups’ LFP recordings. Metyrapone treatment restored the ability of the mother reunion in the rSSP to suppress high frequency oscillations in adversity-reared pups [Fig. 6C; two-way ANOVA, rearing x frequency; main effect of frequency (*F*(3,72)=5.191, *p*=0.003)]. In addition, *post hoc* analyses showed that metyrapone treatment normalized effects of reunion on beta and gamma power to control levels [beta: Control Saline (0.659 ± 0.059) vs. Adversity Saline (1.059 ± 0.101), *p*=0.019, Control Saline vs. Adversity Met (0.705 ± 0.106), *p*=0.648; gamma: Control Saline (0.655 ± 0.034) vs. Adversity Saline (1.061 ± 0.057), *p*=0.035, Control Saline vs. Adversity Met (0.769 ± 0.065), *p*=0.588]. Finally, metyrapone was associated with decreased theta-gamma coupling in maltreated pups compared to saline– treated pups (Fig. 6D), suggesting this treatment rescued control-like cortical phase-locking patterns (X^2^(1,11)=4.412, *p*=0.018, relative risk=0.24).

## Discussion

Here, using a clinically-informed approach, we demonstrate that our novel rat pup rSSP aligns with the human infant SSP to show that adversity rearing experience induces atypical attachment behaviors across species. Using rodents to go beyond the correlational limitations of the human SSP, we capitalize on the power of animal models through randomized assignment, measuring localized brain activity and behavior during experimental treatment/testing and direct assessment of causation. While we acknowledge limitations of using an animal model to understand human behavior, we suggest attachment is particularly well-suited for cross species translation since it is a phylogenetically preserved behavioral system present across altricial species, ranging from birds to mammals^29-31^. Indeed, human attachment theory was based, in part, on animal research^32^, including imprinting in birds^33,34^ and the importance of the emotional bond between the caregiver and infant^35^.

Our data suggest the SSP appears valid across species and represents a translational bridge and access to decades of important and insightful developmental research. Since its inception in 1969^10^, the SSP has been the gold-standard tool for classifying attachment security with strong predictive value for future mental health wellbeing; most notably, disorganized attachment has been shown to be a significant risk factor for later socio-emotional difficulties, including poor stress-management, and future psychopathology^2,3,36^. Our ability to induce aberrant attachment with maltreatment is consistent with the literature, where maltreatment has been shown to statistically mediate the association between poor caregiving and increased risk for later psychopathology and altered neural functioning^24,36-39^. We suggest the present research advances our understanding by identifying causal neurophysiological mechanisms underlying the development and expression of attachment quality.

In typically-reared pups, the mere presence of the mother and maternal behaviors regulate pup cortical LFP oscillations^20^. Translating across species, this regulation by the mother is likely part of a more global regulatory system by which the mother regulates the immature infant’s homeostasis prior to the development of self-regulation^40,41^. This phenomenon is well-documented in children as critical for supporting the child’s response to the world by promoting exploration, as well as across species for decreasing pain/fear, with a critical role for stress hormones^42,43^. The present work suggests this regulation is attenuated in adversity-reared pups during the rSSP as indicated by LFP and behavior towards the mother. Related research on infant rodents’ neurobehavioral response to the mother suggests the neural representation of the mother is reduced^44^, perhaps linked to the internal representation of the mother discussed in attachment theory. While children’s maternal representation engages all sensory systems, pups rely primarily on their mother’s odor^45,46^ and our adversity-reared pups express a less robust preference for the mother and less robust neural responses to the maternal odor than control pups, although the mother remains an attachment figure^47,48^.

To better understand how adversity-rearing caused aberrant rSSP, we recorded pups neurobehavioral responses to the mother during adversity rearing itself. We showed that during the adversity experience pup cortical LFP response to maternal care was blunted. Unexpectedly, this blunting was significantly more salient in response to nurturing maternal behaviors, compared to rough handling by the mother. Specifically, recording pup oscillations during adversity revealed that control and adversity-reared infants showed similar cortical LFP responses to stable states such as non-nursing and pups separated from the mother, while LFP reactivity to *nurturing* behaviors (milk ejection, grooming) was dramatically blunted during adversity bouts (summarized in Figure 7).

Blocking stress hormone synthesis via metyrapone administration restored pup behavior, maternal regulation of LFP power, and cross-frequency coupling patterns to control levels. While the broad neurobehavioral scope of stress hormones to repair brain and behavior may seem surprising, increasing the child’s stress to uncover hidden deficits in attachment was one of Mary Ainsworth’s goals as she developed the SSP^10^. Led by developmental psychology research, animal models have also shown that increasing stress hormones in early life prematurely engages the amygdala in pup social behavior, which provides a “brake” on social behavior^49-51^. Our approach of assessing pups as the maltreatment is ongoing, which naturally produced an increase in pups’ stress hormone levels (present results and^51^), has uncovered that this hormone increase appears causal in the neurobehavioral deficits associated with maltreatment. Cortical and limbic system activity, including gamma oscillations, have previously been shown to be stress-and corticosterone -dependent, even in developing animals^52-55^. The results here demonstrate that these hormonal effects interact with ongoing social and sensory stimulation.

While infant cortical oscillations are well-documented to guide brain development^56-58^, the present results show that the observed perturbations in LFP rhythms are influenced by pup stress hormones and are linked with immediate pup behaviors. Gamma oscillations are a basic mechanism promoting neural synchrony, synaptic plasticity, and information flow. Thus, while these LFP gamma-related outcomes may be involved in expression of the specific behavioral outcomes we quantify here, their modification in itself, especially during early development, is of great importance in understanding the myriad effects of early trauma. Disruption of normal maternal-modulation of gamma oscillations by early trauma could have multiple, long lasting effects regardless of whether they impact specific ongoing behaviors. Indeed, external events and internal state are well-documented to modulate neural oscillations and affect functions including circuit excitability, information flow, and synaptic plasticity^58-62^. For example, theta band oscillation power in cortex and other regions can be modulated during behavioral exploration and are critical for synaptic plasticity, memory, and anxiety-related behaviors in adult humans and animal models^61,63,64^. Furthermore, they are specifically related to childhood performance and predictive of later life outcome^4,65^. Similarly, cortical gamma band oscillations appear to be involved in computations necessary for memory and attention^66,67^. Generation of different frequency band oscillations occurs via different mechanisms; however, cross-frequency coupling between different bands can occur and greatly contributes to information encoding^68,69^. Our results further demonstrate that maternal behavior modulates these cortical LFP oscillations in a specific, behaviorally-dependent fashion^20,70^. However, adversity-rearing modifies pup cortical response to these behavioral events at a developmentally critical time of attachment and cortical development^71,72^.

Theta-gamma coupling, or modulation of gamma oscillations by theta events, is sensitive to developmental events^5,26,73,74^, has been associated with information processing^21,26,75^, and abnormal coupling is diagnostic of neuropsychiatric disorders^76^. Unexpectedly, we observed that adversity-reared pups exhibit more reliable theta-gamma coupling during SSP reunion with the mother than control pups. This same pattern of increased coupling was observed during bouts of adversity rearing compared to controls. This stands in contrast to decreased patterns of theta-gamma coupling in neglected children^4^, not maltreated. However, both abnormally reduced^77^ and enhanced^78^ disruptions of normal cortical cross-frequency coupling are associated with circuit dysfunction and pathology. Furthermore, it is important to note that the timing of assessment necessarily occurs *after* early-life trauma in children, whereas our results reflect immediate changes occurring during/immediately after the adversity. Taken together, increased levels of cross-frequency coupling associated with maltreatment may reflect another example of altered processing of the maternal cue.

Importantly, as shown in Table 1, we did not observe changes in the quality or frequency of nurturing interactions (grooming/nursing) when the mother’s bedding was limited compared to controls. However, the pups’ neural response to distinct maternal inputs, licking/grooming, and milk ejection was blunted during adversity. The similar blunting impact across these inputs is remarkable in light of the fact that these are very different modalities that engage disparate neural networks and neurotransmitter systems. However, a previous study from the lab showed that maternal grooming and milk ejection both appear to regulate pup high-frequency oscillations through a noradrenergic mechanism, as propranolol administration to pups blunted the impact of these maternal behaviors on high frequency oscillations^20^. Although the processing of maternal stimuli is complex and is mediated by multiple systems, this suggests a similar noradrenergic mechanism may mediate the effects of stress on pup processing of these cues. Indeed, corticosterone effects on amygdala plasticity are known to be noradrenaline-dependent^79,80^ and amygdala-cortical pathways involved in processing of the maternal cue are impacted by maternal maltreatment ^81^, suggesting the noradrenergic-HPA interface may provide a useful starting point for understanding the mechanisms for the observed effects. Overall, these data suggest altered processing of the maternal cue, which is supported by other studies showing degraded neural processing of maternal inputs following trauma^44,48,82^.

In conclusion, the present results support the view that sensory stimulation provided by the mother has privileged access to both the immediate regulation of brain and behavior, as well as programming of later-life neurobehavioral function. In our animal model, both adversity-reared and control-reared pups approached and interacted with the mother normally, while the SSP uncovered aberrant attachment to the mother, similar to maltreated and high-risk children^83,84^. The infant’s robust attachment system, which develops regardless of the quality of care received, ensures the infant strives to maintain proximity with the caregiver under conditions of threat, even if the caregiver is abusive. However, while attachment is preserved, its poor quality is associated with pups’ aberrant processing of the sensory stimulation embedded in interactions with the mother, including reduced regulation of pups’ brain and behavior. This information derived from animal models is critical for understanding the roots of pathology induced by early life maltreatment and its causal link to later-life pathology expressed as reactive attachment disorder and other affective disorders.

## Supporting information

Supplementary Methods and Results

## Acknowledgements

The authors would like to thank the following funding sources: NIMH-F32MH112232 (MO), BBRF NARSAD Young Investigator Award (MO), NIH-R37HD083217 (RMS), NIAAA-R01AA023181 (DAW), NIH-R01MH074374 (MD), NIH-R01MH091864 (NT), NYU undergraduate DURF & Wasserman awards (ET), and Gålö-Stiftenlsen (AB). Author contributions: M.O., M.D., R.M.S. and D.W. designed research; M.O., E.T. E.S., T.L., R.M.S., M.D. and D.W. performed research; M.O., E.T., A.B., E.S., K.H., R.M.S., M.D. and D.W. analyzed data; and M.O., E.T., A.B, K.H., R.M.S., N.T., M.D. and D.W. wrote the paper. Thanks to Kira Wood for conducting infant rearing treatments and producing the “stranger” mothers and thanks to Joyce Woo for original illustrations.

## Data Availability Statement

The data that support the findings of this study are available from the corresponding author upon request. The source data underlying Figs. 3-6 and Supplementary Figs. 1-3 are provided as a Source Data file.

