## Supplementary Methods and Results for "Adverse caregiving in infancy blunts neural processing of the mother: Translating across species"

**Supplementary Material**

**Table S1.**

| Rodent developmental trajectories following adverse rearing during infancy | | | |
|---|---|---|---|
| | **Peri-weaning (PN18-23)** | **Adolescence (PN45)** | **Adulthood (>PN60)** |
| **Response to threat** | Increased[1] | Decreased[1,2] | Increased[1] |
| **Social behavior** | Decreased[3] | Decreased[4] | Decreased[4] |
| **Depressive-like behavior** | -- | Increased[3] | Increased[3,5] |
| **Amygdala engagement** | Increased[1,6] | Decreased[1] | Increased[1] |
| **PFC engagement** | None[2] | Increased[1] | Increased[2] |

**Table S1. Long-term neurobehavioral outcomes in threat and social processing following early-life adversity.**

**Table S2.**

| **Maternal Behavior (% total time)** | **Scored during undisturbed activity** | | | | **Scored when returned to nest** | |
|---|---|---|---|---|---|---|
| | A *Data also in Table 1* **Control – No Injection PN8-12** | **LB – No Injection PN8-12** | B **Control – No Injection PN10-14** | **LB- No Injection PN10-14** | C **Control – Injected PN8-12** | **LB – Injected PN8-12** |
| In nest | 54.89 ± 11.63 | 67.6 ± 8.83 | 70.04 ± 4.06 | 68.83 ± 9.09 | 72.58 ± 6.69 | 70.2 ± 7.9 |
| Nursing | 47.52 ± 4.66 | 58.51 ± 9.99 | 53.05 ± 8.91 | 65.83 ± 7.24 | 64.06 ± 7.89 | 58.73 ± 5.67 |
| Milk ejection | 0.59 ± 0.28 | 0.43 ± 0.08 | 0.87 ± 0.14 | 1.02 ± 0.29 | 0.5 ± 0.19 | 0.65 ± 0.06 |
| Rough Handling | 0.68 ± 0.28 | **3.00 ± 0.91** | 0.23 ± 0.11 | **2.09 ± 0.70** | 0.48 ± 0.09 | **3.86 ± 1.43** |
| Grooming | 4.72 ± 3.46 | 5.71 ± 3.91 | 0.65 ± 0.23 | 0.75 ± 0.32 | 10.38 ± 2.6 | 8.13 ± 3.59 |

**Table S2. Maternal care across development and experimental conditions.** Decreasing bedding materials from 4000ml to 100ml produces robust and immediate changes in maternal behavior (A). The relative frequency of rough handling (dragging and stepping on pups) is more frequent in the maltreating mothers, behaviors which typically occur as the mother enters and leaves the nest. Maternal care of older pups (B, PN10-14, Experiment 2) shows a similar increase in rough handling compared to the same treatment from PN8-12 (A). In C, when pups received daily IP administration of drugs ("injected") and were returned to the nest after the drug effects have worn off, increased rough handling was maintained in the low-bedding condition. It should be noted that while high levels of maternal care are typical when pups are placed back into the nest, low bedding mothers immediately begin exhibiting enhanced rough handling of pups, indicating the specific differences associated with rearing conditions (Control vs. Adversity-rearing) were maintained. Bold values are significantly different ($p$<0.05) between Adversity-reared and control. Source data are provided in the Source Data File.

**Table S3**

| | **Low-Risk (*n* = 10)** | **High-Risk (*n* = 11)** | **Total (*n* = 21)** | |
|---|---|---|---|---|
| **Child Characteristics** | | | | |
| Child gender, No. (%) | | | | |
| Female | 4 (40.0%) | 4 (36.4%) | 8 (38.1%) | $\chi^2$(1, N = 21) = .03, $p$ > .05 |
| Male | 6 (60.0%) | 7 (63.6%) | 13 (61.9%) | |
| Child age (months) | | | | |
| *Mean* (*SD*) | 19.47 (5.48) | 20.19 (5.96) | 19.85 (5.61) | $t$(19) = -.29, $p$ > .05 |
| *Range* | 12.12 – 26.94 | 12.22 – 28.32 | 12.12 – 28.32 | |
| Child race/ethnicity, No. (%) | | | | |
| African American | 10 (100%) | 5 (45.5%) | 15 (71.4%) | $\chi^2$(3, N = 21) = 7.64, $p$ > .05 |
| Caucasian | 0 (0%) | 1 (9.1%) | 1 (4.8%) | |
| Hispanic | 0 (0%) | 2 (18.2%) | 2 (9.5%) | |
| More than one race | 0 (0%) | 3 (27.3%) | 3 (14.3%) | |
| **Parent Characteristics** | | | | |
| Parent gender, No. (%) | | | | |
| Female | 9 (90.0%) | 11 (100%) | 20 (95.2%) | $\chi^2$(1, N = 21) = 1.56, $p$ > .05 |
| Male | 1 (10.0%) | 0 (0%) | 1 (4.8%) | |
| Parent age (years) | | | | |
| *Mean* (*SD*) | 30.43 (6.27) | 25.29 (8.03) | 27.74 (7.54) | $t$(19) = 1.62, $p$ > .05 |
| *Range* | 18.78 – 38.71 | 16.14 – 41.00 | 16.14 – 41.00 | |
| Parent race/ethnicity, No. (%) | | | | |
| African American | 9 (90.0%) | 5 (45.5%) | 14 (66.7%) | $\chi^2$(3, N = 21) = 6.11, $p$ > .05 |
| Caucasian | 0 (0%) | 3 (27.3%) | 3 (14.3%) | |
| Hispanic | 0 (0%) | 2 (18.2%) | 2 (9.5%) | |
| More than one race | 1 (10.0%) | 1 (9.1%) | 2 (9.5%) | |
| Parent education, No. (%) | | | | |
| Some high school | 2 (20.0%) | 8 (72.7%) | 10 (47.6%) | $\chi^2$(3, N = 21) = 6.57, $p$ > .05 |
| Completed high school | 6 (60.0%) | 2 (18.2%) | 8 (38.1%) | |
| Some college/trade school | 1 (10.0%) | 1 (9.1%) | 2 (9.5%) | |
| Completed college | 0 (0%) | 0 (0%) | 0 (0%) | |
| More than college | 1 (10.0%) | 0 (0%) | 1 (4.8%) | |
| Parent marital status, No. (%) | | | | |
| Married | 2 (20.0%) | 3 (27.3%) | 5 (23.8%) | |

| | | | | |
|---|---|---|---|---|
| Divorced | 1 (10.0%) | 0 (0%) | 1 (4.8%) | $\chi^2(3, N = 21) = 1.25, p > .05$ |
| Living together | 2 (20.0%) | 2 (18.2%) | 4 (19.0%) | |
| Single | 5 (50.0%) | 6 (54.5%) | 11 (52.4%) | |
| Parent income, No. (%) | | | | |
| Less than $10,000 | 4 (4.0%) | 6 (54.5%) | 10 (47.6%) | $\chi^2(4, N = 21) = 1.36, p > .05$ |
| $10,000 - $19,999 | 2 (20.0%) | 2 (18.2%) | 4 (19.0%) | |
| $20,000 - $29,999 | 2 (20.0%) | 2 (18.2%) | 4 (19.0%) | |
| $30,000 - $39,999 | 1 (10.0%) | 1 (9.1%) | 2 (9.5%) | |
| $40,000 - $59,999 | 1 (10.0%) | 0 (0%) | 1 (4.8%) | |

**Table S3. Child and parent demographic characteristics.**

**Table S4.**

| | **Low-Risk** | **High-Risk** | **Total** |
|---|---|---|---|
| | **(*n* = 10)** | **(*n* = 11)** | **(*n* = 21)** |
| **Child Risk Factors, No. (%)** | | | |
| Low birth weight | 1 (10.0%) | 3 (27.3%) | 4 (19.0%) |
| Prenatal substance exposure | 2 (20.0%) | 5 (45.5%) | 7 (33.3%) |
| Difficult temperament | 0 (0%) | 1 (9.1%) | 1 (4.8%) |
| **Parent Risk Factors, No. (%)** | | | |
| Low income (Income-to-needs ratio < 1) | 4 (40.0%) | 9 (81.8%) | 13 (61.9%) |
| Mental health concerns | 3 (30.0%) | 10 (90.0%) | 13 (61.9%) |
| Low education (Less than high school) | 2 (20.0%) | 8 (72.7%) | 10 (47.6%) |
| Unemployed | 3 (30.0%) | 9 (81.8%) | 12 (57.1%) |
| Criminal justice system involvement | 0 (0%) | 2 (18.2%) | 2 (9.5%) |
| Adolescent parent (Less than 18yo) | 0 (0%) | 7 (63.6%) | 7 (33.3%) |
| Single parent | 6 (60.0%) | 6 (54.5%) | 12 (57.1%) |
| Substance abuse | 2 (20.0%) | 8 (72.7%) | 10 (47.6%) |
| **Instability Risk Factors, No. (%)** | | | |
| Residential (At least one move) | 4 (40.0%) | 9 (81.8%) | 13 (61.9%) |
| Relationship (Status change) | 2 (20.0%) | 7 (63.6%) | 9 (52.9%) |
| Homelessness | 1 (10.0%) | 4 (36.4%) | 5 (23.8%) |
| Child separation (More than 2 weeks) | 0 (0%) | 1 (9.1%) | 1 (4.8%) |
| Other children removed | 1 (10.0%) | 6 (54.5%) | 7 (33.3%) |
| **Total Cumulative Risk Index Score** | | | |
| *Mean* (*SD*) | 3.10 (1.20) | 8.55 (1.92) | 5.95 (3.20) |
| *Range* | 1 – 5 | 6 – 11 | 1 – 11 |

**Table S4. Risk factors and cumulative risk scores by group.**

**Table S5.**

Criteria for coding disorganized behavior in the SSP

I. Sequential Display of Contradictory Behavior Patterns: Child shows behaviors such as avoidance and resistance that represent different attachment strategies sequentially.

II. Simultaneous Display of Contradictory Behavior Patterns: Child shows behaviors such as avoidance and resistance that represent different attachment strategies simultaneously.

III. Undirected, Misdirected, Incomplete, and Interrupted Movements and Expressions: Child directs attachment behavior to stranger, or to no one in particular, for example.

The following four behaviors occurred infrequently and were not considered further.

IV. Stereotypies, Asymmetrical Movements, Mistimed Movements, and Anomalous Postures

V. Freezing, Stilling, and Slowed Movements and Expressions

VI. Direct Indices of Apprehension Regarding the Parent

VII. Direct Indices of Disorganization or Disorientation

**Table S5. Criteria for coding disorganized behavior in the SSP.** Children's disorganized behaviors in the Strange Situation were coded using criteria of Main and Solomon (1990). Using criteria developed by Main and Solomon, the first three behaviors were coded as whether they reached threshold for disorganized behavior.

**Parent sensitivity linking risk with infant disorganized attachment.**

Parenting behavior was assessed through global observational coding of parental sensitivity during a free play activity. This activity was conducted when children were the same age as in the Strange Situation. Parents who displayed high levels of sensitivity tended to respond contingently to their child's cues and adjusted their behavior to the interests and pace of the child. Parents who exhibited low levels of sensitivity failed to respond appropriately to the child's bids, frequently took the lead in the interaction, or appeared detached from the child.

**Figure S1.**

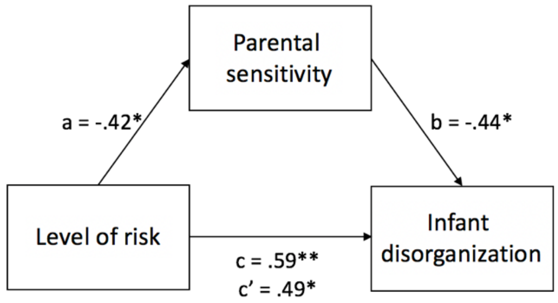


**Figure S1. Model indicating that parent sensitivity is associated with level of risk and infant disorganized behaviors.** Values reflect correlation coefficients. *$p$<0.05, **$p$<0.01

As indicated in Figure S1, parental sensitivity was associated with level of risk and with infant disorganization. When including sensitivity as a co-variate, the association between risk and disorganization was reduced from .59 to .49, suggesting partial mediation. We do not have adequate power to conduct formal mediation analyses, however[7].

**Figure S2.**

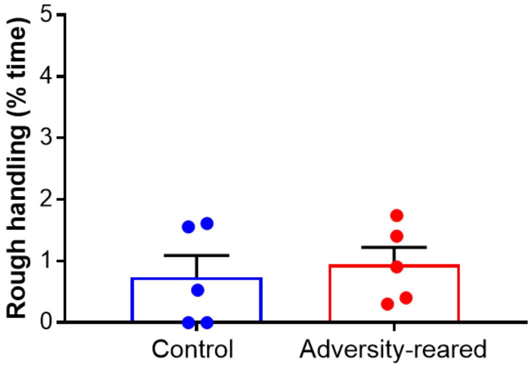


**Figure S2. Within the nest, after adversity-rearing, maternal behavior appears typical.** After litters were brought back to control rearing conditions at the end of the day on PN12 when all mothers were provided with abundant bedding, differences in maternal treatment of pup and pup behavior was not statistically different at PN13-14 ($p$ = 0.71). Source data are provided in the Source Data File.

**Figure S3.**

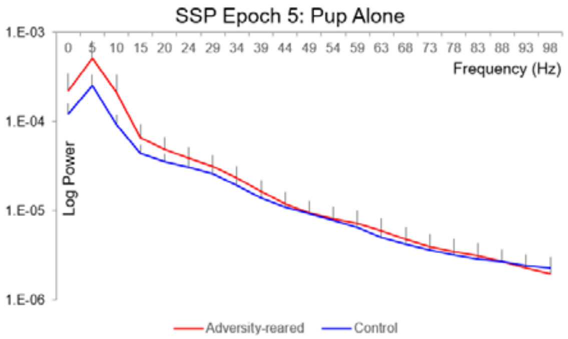


**Figure S3. No group differences in LFP power when pups are alone in SSP.** We did not observe any differences in LFP power in pups that were alone in the SSP (epoch 5), indicating that effects of rearing on LFP power during reunion are not due to baseline differences between groups ($p$>0.05 for all frequency bands). Source data are provided in the Source Data File.

### Supplementary Methods and Materials

#### Subjects

##### *Animals*

Long-Evans rat pups were bred and housed with their mother in polypropylene cages (34x29x17cm) with wood chips and *ad libitum* food and water, in a temperature (20°C), humidity, and light (12h light/dark cycle) controlled room. All procedures pertaining to the use of live rats were approved by the Nathan Kline Institute Institutional Animal Care and Use Committee, and followed National Institutes of Health guidelines. In Experiment 1, a total of 16 pups were used for behavior (1-2 pups per litter from 5 control litters and 5 LB litters; Control 4m 4f, LB 4m 4f); of these, 14 pups were used for LFP analysis. In experiment 2, we used a separate cohort with 1-2 pups from 5 litters during daily 1 hour bouts of control (n = 7, 3m 4f) and LB (n=6, 3m3f). In experiment 3, separate cohorts of pups received either 5 daily injections of metyrapone or saline 90 mins before an hour-long bout of LB/ control (n=6/drug/rearing condition; total = 24, 12m12f) or separate pups received a single drug injection 90 min before SSP following chronic LB/control rearing (n=5-6/drug/condition, total = 22, 11m11f).

***Human Participants***

Parents and their children were referred to the study by Child Protective Services due to risks for maltreatment, such as living homeless, parents having mental health or substance abuse problems, or evidence of prior maltreatment. From this larger sample, two smaller samples were selected for this study. A high-risk group (*n* = 11) was selected that had at least 6 total risk factors and that did not receive specialized training in sensitive parenting. A control group (*n* = 10) was selected that had 5 or fewer total risk factors and did receive specialized training in sensitive parenting. No other criteria were used for selection. Please see Tables S3 for additional information regarding human participant demographic characteristics. At the time children participated in the Strange Situation, they were between the ages of 12.1 and 28.3 months of age (*M* = 19.9, *SD* = 5.6).

**Adversity Experience**

***Modeling infant maltreatment in rodents***

Our Scarcity-Adversity model of low bedding (Adversity-LB) in a cage with a solid floor decreases maternal behaviors and increases rough treatment of pups [4,8]. Rats of both sexes and all ages prefer solid floors [9] with nesting material large enough to build a nest, such as wood shavings or pieces of newspaper. This permits the animal to engage in species specific behaviors such as digging and constructing a nest. Nest building is a particularly potent behavior for mothers, which produces an environment that keeps pups together and provides some shelter. Depriving the mother rat of nesting material is stressful [10] and is well-documented to disrupt typical maternal care, and immediate and enduring disruption of neurobehavioral measures [6,11,12]. This model is similar to another LB model that uses both decreased bedding and a wire-mesh floor, which also effectively disrupts maternal behavior and produces neurobehavioral deficits [13].

In both Experiment 1 and 2, pups lived in either typical nest environments or mothers were given insufficient bedding (1000ml, 1.27cm layer over about half of the floor, compared to the normal 4500ml, 5.08cm layer). This manipulation (adversity-rearing) decreases the mother's ability to construct a nest, resulting in frequent nest building, rough transporting/handling of pups, as well as more licking and less nursing, although pups exhibit normal weight gain after five continuous days of low resource housing[11]. In the present study, we use either a chronic (5 D, PN8-12) treatment (Experiments 1) or brief 1 hr bouts of adversity-rearing once a day (Experiments 2-3) between the pup ages of PN10-14. This short manipulation is validated to modify pup behavior and cortical function [14,15]. As shown in Table S2, maternal care of pups is similar when provided LB across these two ages.

***Risk factors in children.***

*Cumulative risk indices.* Cumulative risk indices were developed across three domains: child, parent, instability, consistent with previous literature [16,17]. Information for the cumulative risk indices were obtained from demographic questionnaires completed by parents, as well as questionnaires regarding parental mental health and child temperament. In addition, a life events interview utilizing a calendar-based method was conducted with the parent. This interview queried the presence of a range of risk factors throughout the child's life. Information from all sources was consolidated. Each risk factor was given a score of zero (0) if absent and a score of one (1) if present, and the total number of risk factors was summed for each child. Risk factors were assessed as present or absent during the child's first two years. Information regarding specific risk factors are included below. See Table S4 for descriptive information regarding risk factors for each group. The high-risk group had a significantly higher cumulative risk total score ($M$ = 8.7, $SD$ = 1.7) than the low-risk group ($M$ = 2.8, $SD$ = 1.2; $t(19) = -9.15$, $p < .001$).

***Child cumulative risk.***

*Child low birth weight.* Parents reported the child's birth weight on the demographic questionnaires. Children who were reported to have a birth weight greater than 2500 gm were given a score of zero (0), whereas those reported to have a birth weight less than 2500 gm were given a score of one (1). Low birth weight (LBW) is defined by the World Health Organization (WHO) as weight at birth less than 2500 gm[18].

*Prenatal substance or alcohol exposure.* Parents reported prenatal alcohol or substance use on demographic questionnaires. In addition, parents were asked about their alcohol and substance use on the life events interview and when they learned that they were pregnant. Children whose parents reported using no alcohol or substances during pregnancy were given a score of zero (0), whereas those whose parents reported using alcohol or substances during pregnancy were given a score of one (1).

*Difficult temperament.* Parents completed the Infant Behavior Questionnaire - Revised (IBQ-R[19]), which is designed to assess temperament in young children. Two scales were used to assess children's temperament: the Distress to Limitations scale, which measures the child's reactions to limitations such as delays in feeding and being placed in a confining position such as a car seat, and the Soothability scale (reversed scored), which assesses the child's reduction of fussing, crying, or distress when the parent uses soothing techniques. Children whose parents scored them less than one standard deviation above the average were given a score of zero (0), whereas children whose parents scored them more than one standard deviation above the average were given a score of one (1).

***Parent cumulative risk.***

*Income.* On the demographic questionnaires, the parent reported yearly family income from all sources (e.g., employment, child support, Temporary Assistance for Needy Families [TANF], etc.). The number of family members living in the residence was also reported on the demographic questionnaires and through the life events interview. An income-to-needs ratio was calculated using this information based on the poverty guidelines which updated periodically in the Federal Register by the U.S. Department of Health and Human Services under the authority of 42 U.S.C. 9902. Parents who reported a ratio greater than 1.0 (above the poverty line) were given a score of zero (0), while those who reported a ratio lower than 1.0 (below the poverty line), were given a score of one (1).

*Parent mental health.* Parent mental health was measured using the Psychiatric Diagnostic Screening Questionnaire (PDSQ), a 125 item self-report checklist designed to assess the presence of psychopathological symptoms [20]. Parents who reported symptoms that exceeded the “clinical cutoff” score for depression, anxiety, or posttraumatic stress disorder on the PDSQ or who reported significant mental health concerns on a life events interview were given a score of one (1). Those who did not report significant mental health concerns on the PDSQ or interview were given a score of zero (0).

*Parent education.* Parents completed demographic information forms listing their education level. Parents who completed high school or beyond (or obtained a GED) were given a score of zero (0). Those who did not complete high school were given a score of one (1).

*Parent employment.* Demographic forms were completed by parents and the life events interview assessed periods of employment. Parents who were employed during the relevant time period were given a score of zero (0), whereas those who were not employed were given a score of one (1).

*Parent criminal justice involvement.* The life events interview and the demographic questionnaires inquired about parental involvement with the criminal justice system. Parents with no criminal justice system involvement during the relevant time period were given a score of zero (0), whereas those with criminal justice system involvement during the time period were given a score of one (1).

*Age first became parent.* Demographic questionnaires completed by parents at the first time point inquired about the parent’s date of birth, as well as the dates of birth of all of their children. Based on this information, the age at which parent first gave birth was calculated.

Parents who were 18 years or older when they first became a parent were given a score of zero (0), whereas those who were 17 years old or younger when they first became a parent were given a score of one (1).

*Single parent.* Information regarding the parent's marital and relationship status was collected through demographic questionnaires and interview questions regarding the presence of a partner during the time period. Parents who reported having a partner for 50% or more of the relevant time period were given a score of zero (0), whereas those who reported not having a partner for more than 50% of the time period were given a score of one (1).

*Parent substance abuse.* The demographic questionnaire given at the first time period assessed for substance abuse, and parents were also asked about a history of substance abuse during the life events interview. Parents who reported no substance abuse during the relevant time period were given a score of zero (0), whereas those who reported substance abuse during the time period were given a score of one (1).

***Instability cumulative risk.***

*Residential moves.* During the life events interview the parent described each residence since the child was born and when residential moves occurred. If no residential moves occurred during the relevant time period, the parent was given a score of zero (0). Parents who reported residential moves occurring during the relevant time period (when the child was 0 to 2 years old) were given a score of one (1).

*Changes in romantic partners.* Parents reviewed their romantic relationships during the life events interview, including periods when relationships began or ended. If the parent reported no changes in romantic relationships during the relevant time period, the parent was given a score of zero (0). Parents who reported the beginning or end of a romantic relationship during the relevant time period were given a score of one (1).

*Homelessness.* Information regarding homelessness was gathered from the initial referral information, as well as through the interview utilizing a calendar method. Parents who reported no homelessness during the relevant time period were given a score of zero (0). Those who reported being homeless at any time during the relevant time period were given a score of one (1).

*Significant separations between child and parent.* Parents described any significant separations (for one month or more) between child and parent in the life events interview. Parents who reported no significant separations during the relevant time period were given a score of zero (0), whereas those that reported a significant separation during the time period were given a score of one (1).

*Removal of other children by CPS.* Parents reported on the removal of other children by CPS on the first demographic form and also in the life events interview. Parents who reported no children removed by CPS were given a score of zero (0), whereas parents who reported other children removed by CPS were given a score of one (1).

**Additional information on the Strange Situation Procedure in children (SSP)**

At young ages, separations from the parent are stressful, and therefore the way that the infant uses the parent for comfort at the time of reunion has proven diagnostic for classifying infant's attachment representations within the range of typically developing children (i.e. secure or insecure) vs. attachment representations associated with later life pathology (i.e. disorganized)[21,22]. The procedure used is the Strange Situation Procedure (SSP) designed by Ainsworth and colleagues[23] originally to identify normative patterns of attachment behaviors exhibited by infants and young children. Briefly, the SSP is a series of separations and reunions between parent and infant. The SSP begins with the parent and infant alone in a new environment. After an unfamiliar female enters the room, the parent leaves the child alone with the stranger. The parent and child are then reunited after 3 minutes. The parent subsequently leaves the child alone in the room (without the stranger) for 3 minutes before the stranger's return, and 3 minutes later the parent returns. The separations are highly potent stressors for the infant, with the stress typically increasing for the infant as the stressor epochs ensue. The behavior of the infant is coded primarily during the two reunions. By naturalistically assessing infants' reliance on the parent when they are distressed[22], infants can be classified as having established secure or insecure attachment patterns. Main et al. (1990)[21] later discovered that the same paradigm could also identify atypical attachment patterns, known as 'disorganized' patterns, that were significantly more common for infants who had been maltreated by their parent. These aberrant attachment patterns were the behaviors of interest in the current study of maltreatment. Infants in the current study were administered the SSP, which typically takes about 24 minutes and includes two separations from, and subsequent reunions with, the parent. Attachment behaviors (proximity seeking, contact maintenance, avoidance, resistance) were coded during reunion episodes. Disorganized attachment behaviors were coded throughout all episodes, with an emphasis on reunion episodes.

***SSP- Humans***

Children in the SSP reunion were classified as secure if they sought out contact with the parent directly and were soothed by the parent, as avoidant if they turned away from the parent or failed to look to the parent for reassurance, and as resistant if they were not soothable despite moving toward the parent. In addition, children were classified as disorganized using criteria specified by Main and Solomon[21]. Children were classified as disorganized if they met the threshold for disorganized behaviors, which included: simultaneous display of contradictory behaviors, sequential display of contradictory behaviors, freezing or stilling, misdirected attachment cues (e.g., approached stranger when distressed), stereotypies or anomalous postures in the parent's presence, direct indices of apprehension regarding the parent (e.g., fearful expression when the parent returns), or direct indices of disorganization or disorientation (e.g., rapid changes in affect, disoriented wandering). Inter-rater reliability was excellent, with the two coders agreeing on 85% of the videos for four-way classifications (k = .74), 92% of secure–insecure classifications, and 87% for organized–disorganized classifications (k = .76). Disagreements were resolved by conference.

**Parent sensitivity.** Parenting behavior was assessed through global coding of parental sensitivity during a free play activity conducted at the same time as the Strange Situation. During this task, parents were instructed to play with their child as they normally would, and were provided with a series of toys that varied based on the child's age. For children under 18 months, children were placed in a high-chair and given a set of three toys (i.e., squeaky toy, rattle, stacking cups). Parents were instructed to interact first at a distance of approximately 3 feet away from the child (without touching the toys) for 2 minutes and then at whatever distance from the child they liked (allowed to touch the toys) for an additional 7 minutes. For children older than 18 months, parents were provided with a set of blocks and asked to play with their children for 7 minutes.

Video-recorded play interactions were coded using a global 5-point scale of sensitivity, adapted from the Observational Record of the Caregiving Environment [24,25]. The sensitivity scale assessed the parent's ability to follow the child's lead by responding appropriately to the child's signals. Parents who displayed high levels of sensitivity tended to respond contingently to their child's cues, and adjusted their behavior to the interests and pace of the child. Parents who exhibited low levels of sensitivity failed to respond appropriately to the child's bids, frequently took the lead in the interaction, or appeared detached from the child. Coders who were blind to study condition, intervention session, date of collection, and study hypotheses were trained to reliability by achieving at least a .75 correlation with a master coder on a reliability set of 10 videos. Inter-rater reliability was further assessed by randomly selecting 20 percent of videotapes for double-coding. A one-way random effects intra-class correlation revealed an ICC of .79.

#### *SSP- Rodents*

In Experiment 1, the SSP was modified to accommodate infant rodents. After 5 days of either control rearing or chronic Scarcity-Adversity LB rearing from PN8-12, pups were tested in a modified SSP apparatus (polypropylene cage with 2000ml bedding, [34x29x17cm]). In the rodent paradigm, a “stranger” rat mother was produced by changing the maternal odor via the diet (Tekland). Rat pups cannot see or hear until they are about PN15 and are hence identifying their mother by the maternal odor, which is dependent on the mother’s gut bacteria and is learned by pups [26,27], rather than a pheromone. Since a laboratory typically feeds the mothers the same diet, all mothers have the same maternal odor. This has led to some incorrectly assuming that pups cannot identify their mother; however, when mothers are individual scented, pups readily discriminate between their own mother and a stranger mother fed a different diet.

Each pup completed one session of the experiment during one day. With the mother and stranger anesthetized with urethane, the rat pup (PN13-14) was placed in a plastic chamber [34x29x17cm] for the duration of the procedure, which lasted approximately 35 minutes. Urethane anesthesia inhibits milk letdown without odor confounds associated with isoflurane and using anesthetized dams prevents behavioral interference from the mothers in the assay of pup attachment behavior. Furthermore, pups cannot see at the ages tested and are primarily guided by the maternal odor; the presence of this odor alone is sufficient to drive typical pup attachment responses [10]. The SSP for the rodent model was divided into seven 5-minute episodes, or epochs: 1) Biological mom (M) and pup (P) are alone in the cage; 2) Stranger (S) is added to cage; 3) M is removed and S and P are in the cage together; 4) S is removed and M and P are in the cage together; 5) M is removed and P is alone in the cage; 6) S is added back to the cage; 7) S is removed and M is added back to the cage. Pup behavior was videotaped during the entire procedure.

**Rodent Behavioral Analysis**

Each experiment was recorded and scored by two highly trained, independent raters and analyzed offline. In SSP experiments 1 and 3, raters were blinded to the previous rearing condition of pups. We scored mother and pup interactive behavior using BORIS ethology software [28] based on video recordings.

For scoring homecage observations and validating rough handling by the mother, pups were videotaped three times a week and data analyzed by Ethovision (Noldus) for automatic scoring of activity in two arenas, outside the nest and in the nest, which is validated to capture the higher activity levels within the nest of the abusive mother [29] and highlights LB mother's higher activity score in the nest compared to controls. Videos are also hand scored using Boris software for the following behaviors: time spent in nest/outside of nest, rough handling of pups (stepping on pups, dragging pups), nursing, milk ejections, grooming pups, scattered litter, mother eating/drinking/self-grooming, and sleeping. Hour-long videos from control and LB litters were segmented into 5 minute bins and the percent of bins where given behaviors were observed was calculated.

***Strange Situation Procedure Analysis***

In Experiment 1, we present data from the last epoch. The pup behaviors measured were categorized as typical (pup crawls to mother, pup probing mother, pup sleeping/laying on mother) or atypical (pup sleeping/laying alone, pup sleeping behind the mother).

***Homecage observations***

We present our results within the sequence of typical mother-infant interactions within the homecage, which is summarized in Figure 7. Within the age range used here, the mother typically remains with the pups but will briefly leave the nest to take care of her own biological needs. Pups typically remain in the nest. When the mother returns to the nest, she begins to interact with pups, focusing on grooming pups and adjusting the nest; in the Scarcity-Adversity model, rough handling of pups typically occurs at this point. Next, pups attach to the mother's nipples, and the mother typically briefly continues to groom but quickly settles down to non-nutritively nurse. Periodically, the mother gives pups a brief pulse of milk (milk ejection), which produces a stretch reflex in pups (tonic elongation of the pup's body followed by increased general activity, lasting a few seconds). While this sequence of behavioral interactions remains constant across Adversity-LB and control rearing, Table 1 illustrates that the relative frequency of rough treatment of pups by the mother is significantly higher when the mother has low bedding (Table 1, $t_{(8)}$ = 2.08, $p$ = 0.037), while nurturing behaviors (nursing, grooming) occurred at control levels (all p's > 0.05) – all typically occurring as the mother enters and leaves the nest. This observation is consistent with previous work from our lab and others [11,12].

In Experiment 2, behaviors were scored for freely-behaving mothers and pups within the nest. In addition to the above-listed behaviors, the behavioral states measured were pup in/out of nest, nursing, milk ejection, and grooming. Abusive behaviors, including dragging pups and rough handling were similarly coded. During nursing, the pup is attached to the mother's nipple, which is typically non-nutritive, although punctuated with periodic milk ejections, which induced a robust stretch reflex in pups (elongation of body for a few seconds). 30s bouts of non-nutritive nursing were identified (>15 examples per pup) and average power per frequency calculated for each pup; this data was used to normalize LFP response during milk ejection, grooming and rough handling. After observing the stretch reflex associated with milk ejection, we measured LFP for ten seconds immediately after the start of this observed behavior. Similarly, we coded for grooming for a ten-second period immediately after the start of observing this behavior. Rough handling LFP was analyzed across bouts ranging from 5-10 secs.

### Electrophysiology

#### *Surgical Procedures*

Pups (PN9-13) were anesthetized and kept unconscious with an isoflurane anesthesia system (E-Z Systems, Palmer, PA) during surgery. Litters were culled to 5 pups on PN1 and 1-2 pups per litter was implanted with a telemetry transmitter. Pups were placed in a stereotaxic apparatus under aseptic conditions. The scalp was reflected and skull dried. A hole was drilled for the recording electrode using coordinates to target the frontal neocortex (~2 mm anterior to bregma, ~2mm laterally over the left hemisphere, ~1.0mm ventral to the surface of the brain) and a hole drilled for the reference electrode over the posterior right hemisphere. A teflon coated 0.18 mm diameter stainless steel electrode was lowered to the desired depth and dental cement was placed over the hole to hold the electrode in place. The electrode was connected to a telemetry pack (ETA-F10 telemetric device, DSI) inserted into the pup subcutaneously on the animal's back. Topical lidocaine hydrochloride jelly (2% Akorn) was applied to the wound and closed with sutures. A small amount of glue (Vetbond) was then placed over each suture. Anti-nail biter liquid was also applied to the area around the sutures to prevent the mother from over-grooming the wound. Prior to waking, each pup was injected with 1cc of 0.9% saline. Upon waking, pups were placed in an incubator for 30 minutes to 1 hour until observed to be fully recovered and mobile, then returned to the mother and continuously observed for several hours to ensure that the mother allowed the pup to nurse. The Sullivan and Wilson labs have extensive experience implanting pups that are behaviorally indistinguishable from non-implanted pups and no evidence of altered maternal behavior towards the instrumented pup is observed[10,30]. Recordings targeted the left frontal cortex to allow consistency with previous work exploring maternal effects on the infant cortical activity [30].

***LFP Recordings***

Neural signals from both telemetry systems were filtered (0.5 to 200 Hz), digitized at 2 kHz with Spike2 software (CED, Inc), and analyzed offline. Spontaneous LFP activity was recorded during the 35 min Strange Situation procedure on one day (Experiment 1) or for 4-5 consecutive days in the freely behaving infants and mothers within the nest (Experiment 2). In Experiment 1, the pup's LFP was recorded over the course of the SSP procedure and analyzed by epoch and across pup behaviors toward the mother and stranger. Experiment 2, the experimental pup and mother were recorded for an hour-long control nesting bout. Then, nesting materials were removed by the experimenter and behavior/LFP recorded for one hour; these phases were counterbalanced. During the recording session, behavioral states of the videotaped pup were noted on the neural trace for off-line analysis. These behavioral states included when the mother entered the nest area, when she left the nest area, when the pup was nursing or attached to the nipple, when the pup received a milk ejection, when the mother roughly handled pups (stepped on/dragged) and when the mother groomed the pup. Following data collection, periods of time during each of these behavioral states could be assessed individually. Recordings within this nest environment lasted for 1 hour and were viewed and recorded via Logitech Webcam Software from a camera positioned over the cage.

***LFP Data analysis***

Fast Fourier Transform (FFT) power analyses were performed on the raw LFP data in intervals taken from sections of each day's neural trace that correlate with a specific behavioral state (described below) to quantify LFP oscillatory power in 2.9 Hz frequency bins from 0–100 Hz (Hanning). Power in the delta (0-5 Hz), theta (5–15 Hz), beta (15-35 Hz) and gamma (35-100 Hz) frequency bands was calculated for each specified window. Behavioral states were noted online by observing the pup and mother behavior. Offline, these notations were used to find and independently assess each behavioral state within a daily recording session. Each animal provided either a single ~30 min recording during the SSP (Experiment 1) or 2-4 hour-long recordings across brief bouts of control bedding or low bedding each day following implantation, and from these recordings, we obtained identical numbers of each behavioral state where available.

***Behavioral states and specified time window of statistical analysis***

Comparisons were made both across-animals and within-animals for each behavioral state (nipple attached, milk ejection, grooming, mom roughly handling pups) or between epochs in the SSP (power across epoch 5 was used as the baseline for normalization). ANOVAs were run on the ratio of LFP power to baseline (30 sec nipple attached, no milk ejections) to test for main effects of behavioral state followed by post hoc analyses to examine differences between specific LFP frequencies. As above, ratio in the power in the theta (5– 15 Hz), beta (15-35 Hz), and gamma (35-100 Hz) frequency bands was calculated for each specified window.

#### *Cross-frequency coupling analysis*

LFPs were filtered into theta (1.5–12 Hz) and gamma (35–100 Hz). Cycles (the nadir of negative troughs) were then detected based on threshold crossing from filtered signals. Phase locking was assessed for each oscillatory frequency band as follows: Experiment 1, assessments during epoch 7 of the SSP (5 min); Experiment 2, assessments during 1 hr bouts of LB and control. MATLAB software was used to determine significance of phase relationship between theta and gamma cycle using Rayleigh statistics for circular uniformity (MATLAB circular statistics package). Comparisons of the proportion of pups showing phase locking were performed using $\chi^2$ ($p < 0.05$). In addition, the mean phase angle and the mean resultant vector length was calculated for each pup in each condition using MATLAB circular toolbox (scripts: circ_mean and circ_r). Population vectors representing the average of the mean phase angle and resultant vector length in each condition were then calculated using MATLAB circular statistics toolbox [31].

#### *Drug Manipulations*

In Experiment 3, separate cohorts of pups received i.p. injections of the corticosterone inhibitor metyrapone HCL (50 mg/kg, Sigma) or an equal volume of 0.9% saline. To block corticosterone changes associated with the SSP following maltreatment, pups that had undergone 5 days of continuous Adversity-LB/control rearing (Experiment 1 timeline) received metyrapone or saline 90 min before SSP testing on PN13-14 and were returned to the homecage before SSP administration. To block corticosterone changes associated with acute maltreatment (Experiment 2 timeline), PN10-14 pups experienced 5 days with 1 hour daily experience with low bedding. Each day, 1 male and 1 female from each litter received an i.p. injection of metyrapone HCL or an equal volume of 0.9% saline and were returned to the homecage. 90 minutes after injection, half of the nursing dams' bedding was reduced from 4500 mL to 100 mL (low bedding), for 1 hour and then control bedding levels returned. Timing of this procedure limited corticosterone reduction effects to within the daily hour-long window of Adversity-LB. Previous work from our lab and others has shown that infant stress increases pup CORT levels and that metyrapone reliably decreases CORT levels by 55-75 percent in pups in this age range[32-36].

***Statistical Analysis***

Depending on the number of groups to be analyzed, behavioral and LFP data were analyzed with Student's t-tests and one- or two-way analysis of variance (ANOVA), followed by post-hoc Bonferroni tests, or $X^2$ analysis and circular statistics (coupling data). Data used for figures are expressed as mean (±SEM) and in all cases, differences were considered significant when $p < 0.05$.
